# Indoleamine Dioxygenase and Tryptophan Dioxygenase Activities are Regulated through Control of Cell Heme Allocation by Nitric Oxide

**DOI:** 10.1101/2022.12.30.522347

**Authors:** Pranjal Biswas, Dennis J. Stuehr

**Author notes:** To whom correspondence should be addressed: Dennis J. Stuehr, Department of Inflammation and Immunity, NC2-103, Lerner Research Institute, Cleveland Clinic, 9500 Euclid Avenue, Cleveland, OH, 44195.

## Abstract

Indoleamine-2, 3-dioxygenase (IDO1) and Tryptophan-2, 3-dioxygenase (TDO) catalyze the conversion of L-tryptophan to N-formyl- kynurenine and thus play primary roles in metabolism, inflammation, and tumor immune surveillance. Because their activities depend on their heme contents which range from 30- 60% heme-saturated in biological settings and go up or down in a dynamic manner, we studied how their heme levels may be impacted by nitric oxide (NO) in mammalian cells. We utilized cells expressing TDO or IDO1 either naturally or via transfection and determined their activities, heme contents, and expression levels as a function of NO exposure. We found NO has a bimodal effect: A narrow range of very low NO exposure promoted cells to allocate heme into TDO and IDO1 and boosted their activities several fold, while beyond this range the NO exposure transitioned to have a negative impact on their heme contents and activities. NO did not alter dioxygenase protein expression levels and its bimodal impact was observed when NO was released by a chemical donor or was generated naturally by immune-stimulated macrophage cells. NO-driven heme allocations to IDO1 and TDO required participation of a GAPDH-heme complex and for IDO1 required chaperone Hsp90 activity. Thus, cells can up- or down-regulate their IDO1 and TDO activities through a bimodal control of heme allocation by NO. This mechanism has important biomedical implications and helps explain why the IDO1 and TDO activities in animals go up and down in response to immune stimulation.

## Introduction

Indoleamine-2, 3-dioxygenase (EC 1.13.11.52; IDO1) and Tryptophan-2, 3-dioxygenase (EC 1.13.11.11; TDO) are heme proteins which catalyze the conversion of tryptophan (Trp) to N-formyl-kynurenine (1,2). The activities of these enzymes directly depend on the content of their ferrous heme, which binds di-oxygen to enable its insertion into the indole ring of L-Trp (3,4). TDO is selective for L-Trp as a substrate while IDO1 accepts a broader range of indole-containing molecules (5,6). Although IDO1 has a monomeric structure and TDO is tetrameric (7), their crystal structures indicate there are similarities in their heme binding environments and substrate binding sites (8), consistent with their having similar catalytic mechanisms (9,10). TDO expression and activity levels are highest in the liver and upregulated by glucocorticoids and L-Trp (11,12). In comparison, IDO1 is constitutively expressed in tissues such as the lung and its expression can be broadly induced by stimuli associated with immune activation and inflammation, including interferon-γ (IFNγ), lipopolysaccharide (LPS), and tumor necrosis factor (TNF) (13-15). Increased IDO1 or TDO activities cause Trp depletion and increased production of Kynurenine (Kyn) and its metabolites, which are bioactive (16-19). In this way, increased IDO1 activity helps down-regulate several inflammatory diseases (20,21), but on the other hand, is also associated with neurologic disorders (22,23), and in cancer cells or in host dendritic cells is associated with suppression of effector T-cell responses toward tumors (24,25). Thus, careful control of the Trp dioxygenase activities is needed and they are targets for pharmacologic intervention (26-28).

We recently reported that maturation of functional IDO1 and TDO requires a GAPDH-dependent heme delivery and in the case of IDO1 also requires heat shock protein 90 (Hsp90) to drive its heme insertion (29). Moreover, we found that in cells under normal growth conditions both IDO1 and TDO were 40-50% heme saturated (29), consistent with earlier reports showing that 30-60% of rodent liver TDO normally exists in a heme-free state in healthy animals (30-32) and that IDO1 is predominantly expressed in its heme-free form in cells (33,34). Older studies also showed that the heme saturation levels of TDO and IDO1 could be dynamically altered: The heme level of TDO could be increased by giving the animals Trp (35,36) or by boosting their heme biosynthesis (31,32). Moreover, these changes could be caused by immune stimulation: Rats injected with the immune stimulant *E. coli* bacterial lipopolysaccharide (LPS) displayed a temporal increase in liver TDO heme content and activity, reaching a maximum 4-6 h post injection and then returning back to or falling below the original baseline after 10 h (37,38). A similar bimodal effect was observed for cellular IDO1 activity in response to various immunologic stimuli (39,40). Although such dynamic regulation of TDO and IDO1 activities is likely to be biomedically relevant, how it occurs upon immune activation is currently unknown.

In researching this topic we noticed that the temporal changes in rat liver TDO heme content and activity following LPS injection were quite similar to the induction of NO synthesis activity that is caused by a similar LPS injection in mice (41). We therefore investigated a possible role for NO in driving these immune-related changes in the activities and heme contents of the two dioxygenases. NO at relatively high levels is already known to block heme insertion into several heme proteins including inducible NO synthase (NOS) (42), endothelial NOS, neuronal NOS, two cytochrome P450 enzymes (CYP 3A4 and 2D6), catalase, and hemoglobin (43). Conversely, lower levels of NO were shown to cause cells to insert heme into the heme-free form of soluble guanylyl cyclase β subunit (sGCβ) and thus trigger assembly of the sGCαβ heterodimer (44), which is the functional NO sensor. In our present study we investigated mammalian cells in culture that either naturally express TDO or IDO1 or did so upon transfection and utilized NO generated either by a well-characterized NO donor compound or by macrophage cells induced to express NOS. We determined IDO1 and TDO activities and heme contents by measuring Kyn production and radiolabel heme incorporation, respectively, and we also investigated possible roles for cellular GAPDH and Hsp90 in the NO-directed processes. Our findings establish that TDO and IDO1 activities in cells are beholden to NO-driven effects on their heme contents. This helps to explain how immune stimulation regulates their activities and reveals a new mechanism by which IDO1 and TDO function may be up- or down-regulated in health and disease.

## Results

### NO has a concentration-dependent, bimodal effect on IDO1 and TDO dioxygenase activities

We first examined how varying NO exposure would impact the activities of TDO and IDO1 in cells. Human cell lines that either constitutively express human TDO (HepG2) or did so upon transfection (HEK293T) were cultured for 12 h in the presence of varying concentrations of the slow-release NO donor NOC-18, whose half-life is reported to be 13 h at pH 7.0 and 37°C (https://www.dojindo.eu.com/store/p/237-NOC-18.aspx) and was calculated to have a half-life of 8 h based on measurements of NO release from NOC-18 under our particular cell culture conditions (Fig. S1). HEK293T cells that had been transfected to express human IDO1 underwent identical treatment. Dioxygenase activities were judged by measuring the product Kyn that had accumulated in the culture fluid over the 12 h period. Figure 1 shows that cells exposed to a low range of NOC-18 concentrations (0.1 to 5 µM) increased their TDO and IDO1 activities between 4 and 6-fold in a concentration-dependent manner. The effect then steadily ebbed such that cells receiving 25 µM NOC-18 displayed near basal levels of activity while those receiving NOC-18 at 50, 75, or 100 µM had activities below the basal levels. Western blot analysis showed that the changes in activity were not due to changes in the levels of TDO or IDO1 protein expression in the cells, which remained similar across the range of NOC-18 exposures (Figs. S2-S4). Thus, IDO1 and TDO when expressed in cells underwent a concentration-dependent bimodal change in their activities in response to NO exposure that was independent of the cell identity or the manner of enzyme expression (natural *vs* transient transfection).

**Figure 1.**
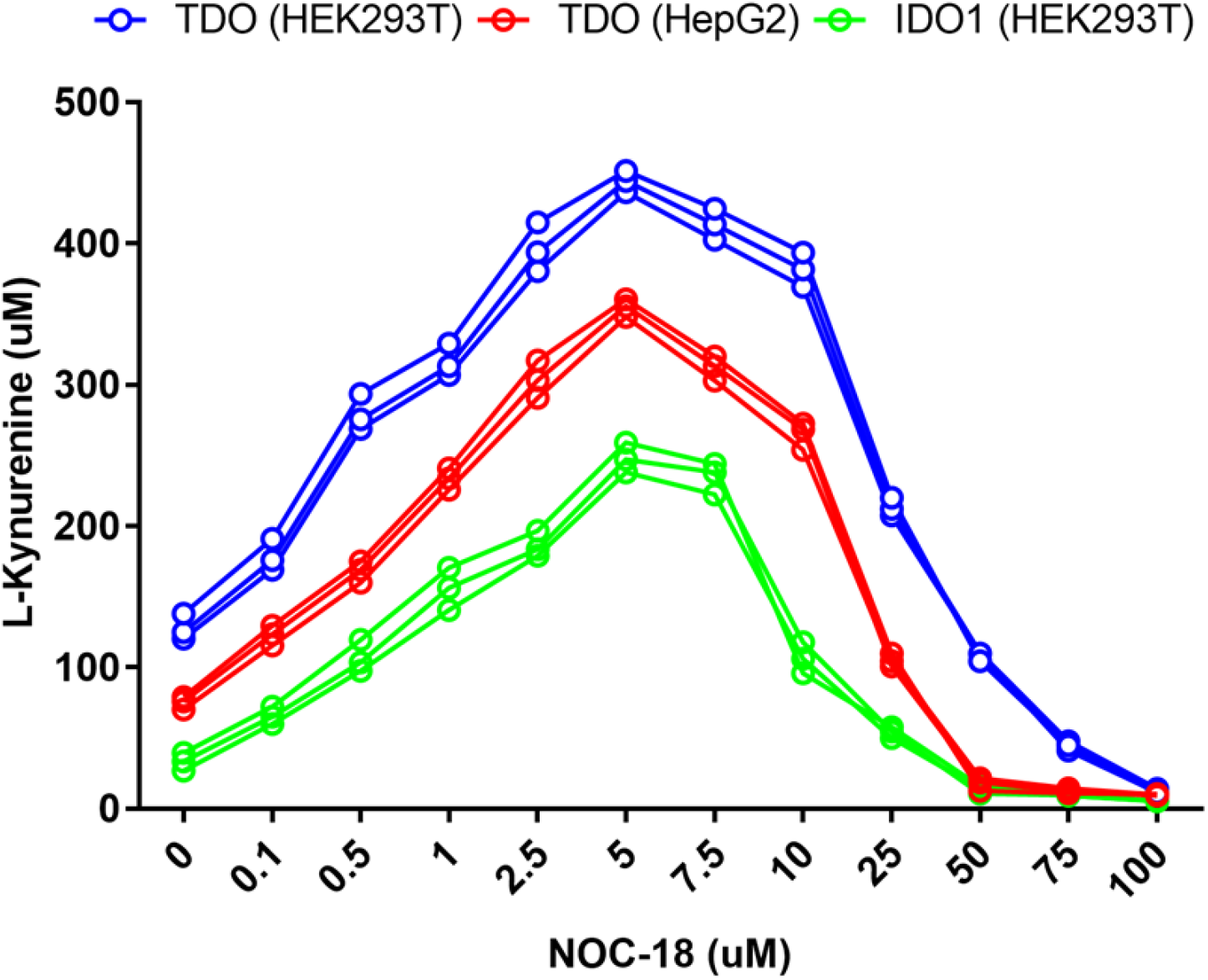
NO regulates cell TDO and IDO1 activities in a concentration-dependent and bimodal manner. The indicated cell lines expressing TDO or IDO1 were cultured 12 h with the indicated concentrations of NOC-18 and product Kyn that accumulated in the media was measured. Data are the mean ± s.d.; n=3 experiments; ***p<0.001, one-way ANOVA.

### The NO effect on dioxygenase activities correlates with an effect on their heme contents

We next examined if the changes in IDO1 and TDO activities were related to changes in their heme contents. Glycine auxotroph Chinese hamster ovary cells (GlyA-CHO) were transfected to express human TDO-FLAG and IDO1-FLAG proteins in ^14^C-Glycine-containing medium for 48 h, after which protein expression was stopped with cycloheximide. Cells were then treated with NOC-18 at different concentrations for 12 h and the effects on the IDO1 and TDO activities & ^14^C-heme contents were determined by measuring Kyn production and by immune-precipitation with anti-FLAG antibody and ^14^C scintillation counting, respectively. Controls included cells not transfected to express the dioxygenases, and transfected cells whose heme biosynthesis was intentionally blocked by inclusion of succinyl acetone (29,43), which enabled us to calculate what percentage of ^14^C counts in the IDO1 or TDO pull downs were due to their ^14^C-heme incorporation *versus* ^14^C-Gly incorporation into the protein chains. Counts due only to ^14^C-Gly in the proteins averaged around 15-20% of the total counts (data not shown) and were subtracted from the total sample counts in each case to obtain the ^14^C-heme specific counts.

Figure 2 panels A and B show that NOC-18 treatment of the GlyA-CHO cells had concentration-dependent and bimodal impacts on the IDO1 and TDO activities that were essentially identical to what we had observed when these enzymes were expressed in the HepG2 and HEK293T cells. Figure 2 panels C and D show that the change in activities correlated directly with changes in the level of ^14^C-heme bound within IDO1 or TDO. Their ^14^C-heme contents rose when cells were given to 5µM NOC-18 to reach an approximate 3-fold increase in ^14^C-heme, and then steadily fell at higher NOC-18 concentrations such that IDO1 and TDO contained only basal levels of ^14^C-heme in cells given 25 µM NOC-18 and contained sub-basal levels in cells given 50 µM NOC-18 or above. TDO or IDO1 protein expression in the cells remained similar across the range of NOC-18 exposures (Figs. S5-S6). Thus, NO-driven changes in IDO1 and TDO heme contents correlated with the NO-driven changes in their catalytic activities.

**Figure 2.**
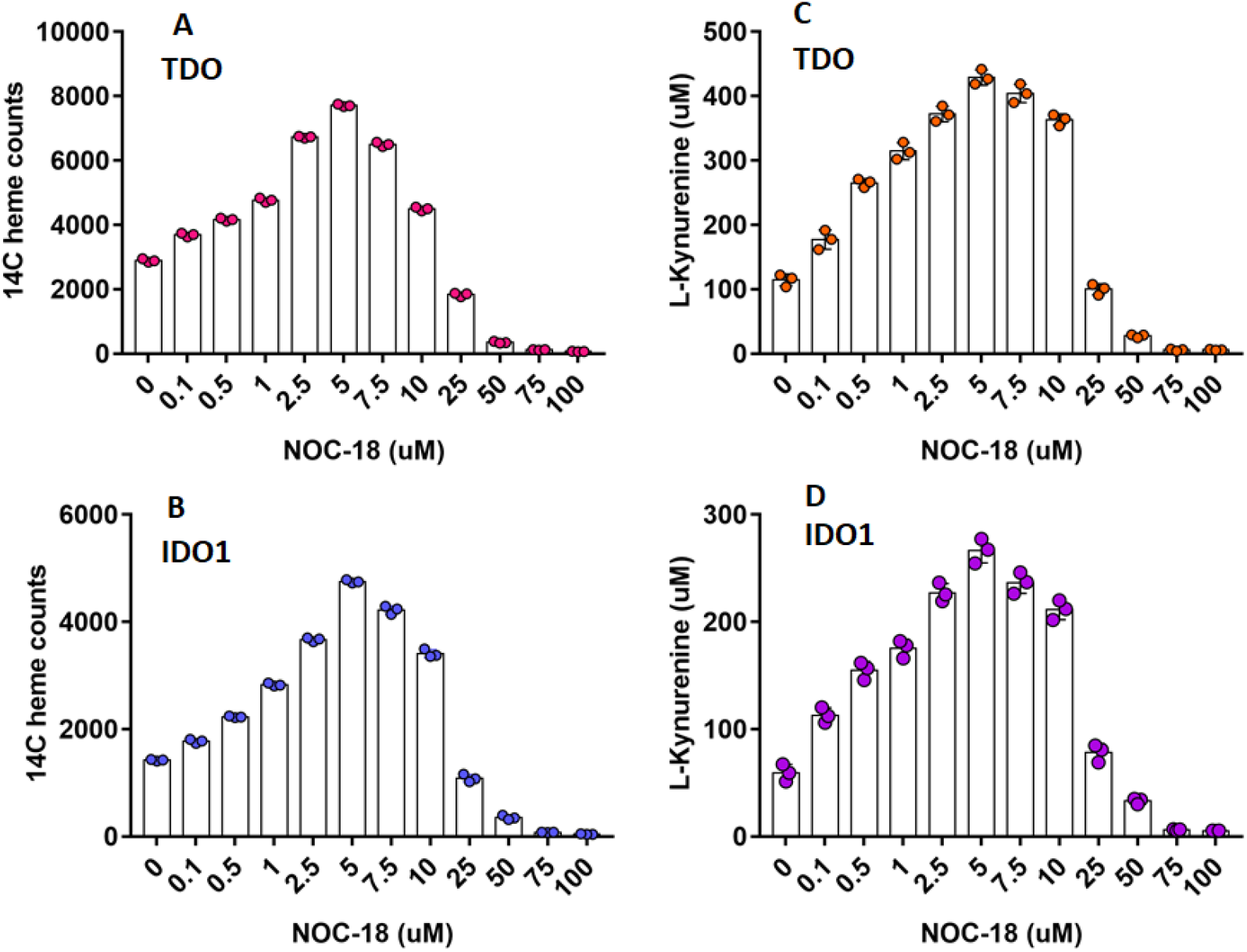
NO regulates the activities of TDO and IDO1 by regulating their heme contents in cells. GlyA-CHO cells were transfected to express TDO-FLAG and IDO1-FLAG in ^14^C-Gly containing media for 48h after which protein expression was stopped with Chx and NOC-18 was added at the indicated concentrations and culture resumed for an additional 12 h, followed by assay of Kyn accumulation and Ab pull down of FLAG-labeled TDO or IDO1 to measure their ^14^C-heme contents. Panels A and B, IDO1 and TDO activities. Panels C and D, IDO1 and TDO ^14^C-heme counts. Data are the mean ± s.d.; n=3 experiments. ***p<0.001, one-way ANOVA.

### NO generated by immune-stimulated cells alters TDO and IDO1 activities and heme contents

To test if NO naturally generated by immune-stimulated cells would have a similar effect as the NO released from NOC-18, we performed a co-culture experiment (Figure 3A) where RAW264.7 macrophage cells that had been cultured in permeable trans-well inserts and treated with *E. coli* lipopolysaccharide (LPS) to induce expression of inducible NOS and NO production were placed above a monolayer of GlyA-CHO cells that expressed either TDO-FLAG or IDO1-FLAG that had ^14^C-heme incorporated due to prior culture of the cells with ^14^C-glycine. The trans-well inserts contained two different quantities of activated RAW264.7 cells (25% and 100% confluent) to create a lower and higher NO production condition, respectively. The co-cultures were analyzed at 0 h or after 6 h had elapsed. Figure 3B shows that nitrite, a NO-derived product, was generated and accumulated in the co-culture fluid after 6 h at levels corresponding to the two different quantities of RAW264.7 cells that were present in the inserts. The GlyA-CHO cells co-cultured for 6 h with the lesser number of activated RAW264.7 cells displayed an increase in their TDO activity (Fig. 3C), while those co-cultured with the higher quantity of activated cells displayed a decrease in TDO activity. These activity changes correlated with a gain or loss in TDO ^14^C-heme level relative to what was initially present in TDO in the cell cultures analyzed at t = 0 h (Fig. 3D). Nearly identical results were obtained in co-culture experiments where expression of IDO1 substituted for TDO in the GlyA-CHO cells (Fig. 4). In replica experiments that used trans-well inserts containing non-activated RAW264.7 macrophages we observed no change in the cell TDO or IDO1 activities or their heme contents after 6 h co-culture (Fig S7). The expression levels of TDO or IDO1 were similar across all cultures and conditions (Fig. S8). Thus, NO from immune-stimulated macrophage cells caused a concentration-dependent bimodal change in cell TDO and IDO1 activities and heme contents that mirrored what we observed for cells exposed to low or high levels of NO released by the donor NOC-18.

**Figure 3.**
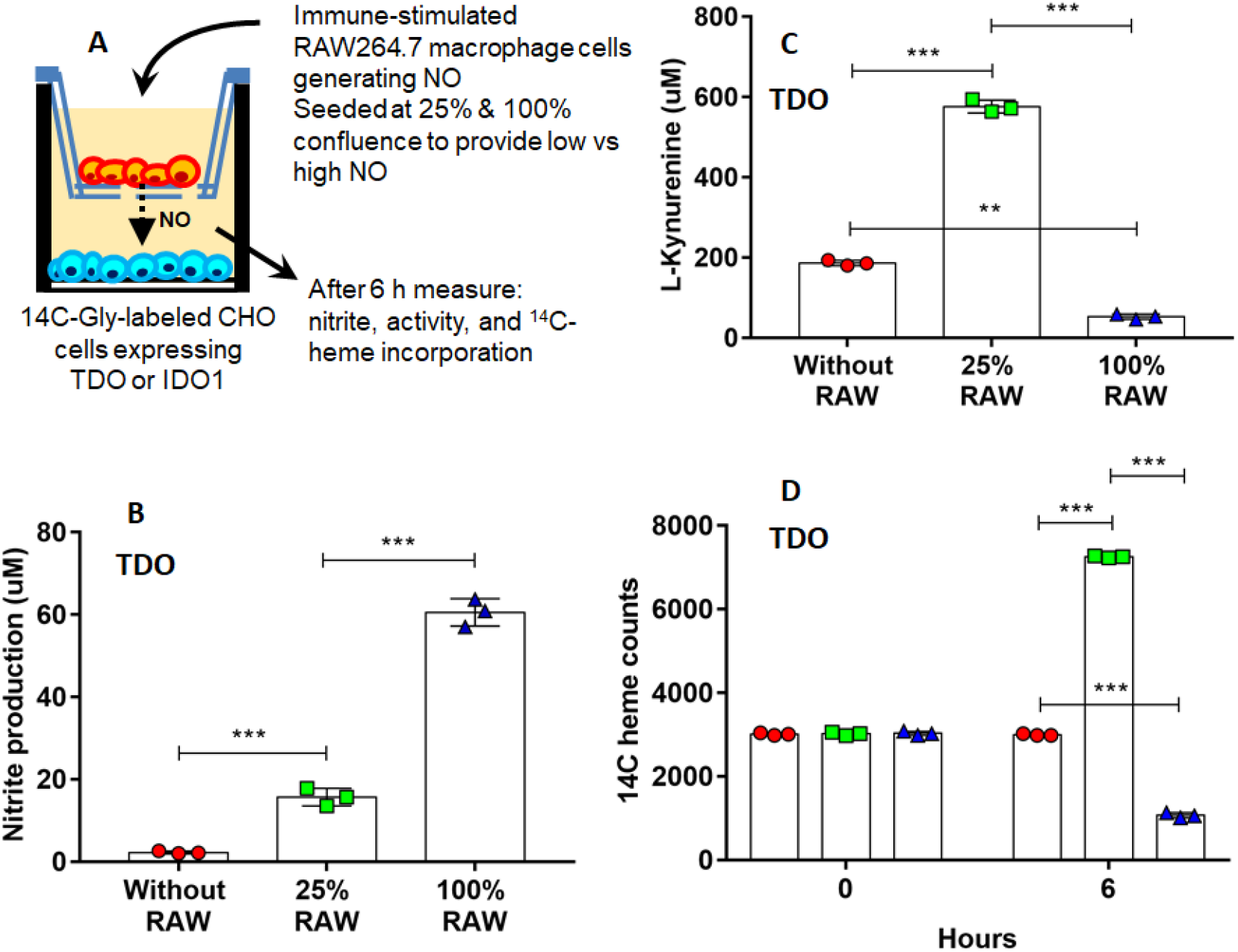
NO released by immune-activated macrophage cells regulates the heme level of TDO expressed in underlying cells. Panel A, trans-well inserts containing RAW264.7 macrophage cells at 0, 25, and 100% confluency that had been activated by bacterial LPS to express NOS and generate NO were placed into culture wells that contained a monolayer of GlyA-CHO cells expressing TDO-FLAG that contained ^14^C-heme incorporated due to prior culture of the cells with ^14^C-Gly. After 6 h of co-culture we measured nitrite (Panel B) and Kyn (Panel C) produced in the cell cultures, and measured ^14^C heme counts in TDO-FLAG pulled down after 0 h and 6 h of co-culture (Panel D). Data are the mean ± s.d. of 3 experiments. ***p<0.001, **p<0.01, one-way ANOVA.

**Figure 4.**
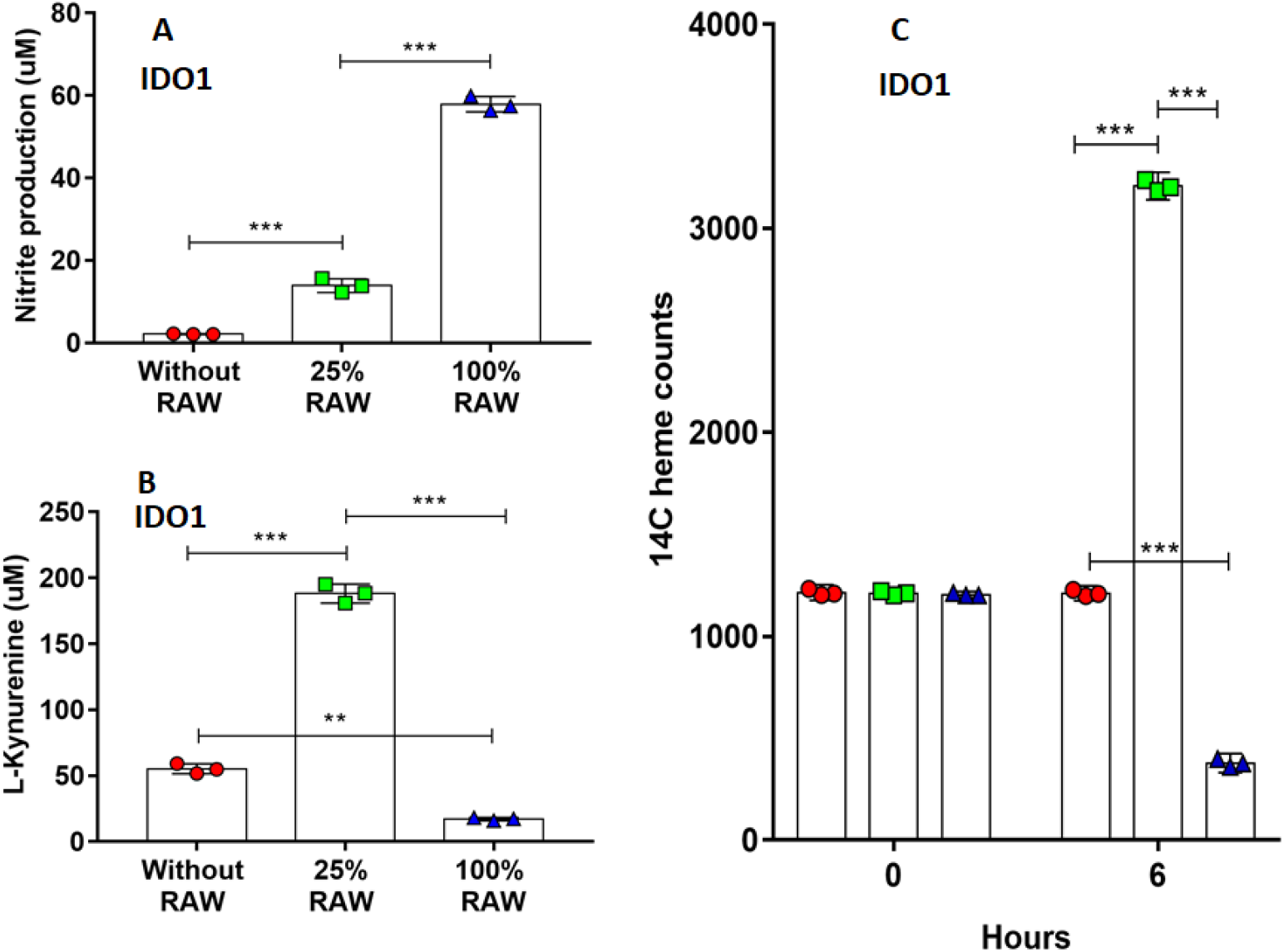
NO released by immune-activated macrophage cells regulates the heme level of IDO1 expressed in underlying cells. Trans-well inserts containing RAW264.7 macrophage cells at 25 and 100% confluency that had been activated by bacterial LPS to express NOS and generate NO were placed into culture wells that contained a monolayer of GlyA-CHO cells expressing IDO1-FLAG that contained ^14^C-heme incorporated due to prior culture of the cells with ^14^C-Gly. After 6 h of co-culture the nitrite (Panel A) and Kyn (Panel B) produced in the cell cultures was measured, and the ^14^C heme counts in TDO-FLAG pulled down after 0 h and 6 h of co-culture were measured (Panel C). Data are the mean ± s.d. of 3 experiments. ***p<0.001, **p<0.01, one-way ANOVA.

### Time course of the NO-driven changes in dioxygenase activities and heme contents

To investigate the kinetics of the NO-driven changes we expressed TDO or IDO1 in ^14^C-Gly-treated GlyA-CHO cells and then exposed them to media alone or either a lower (5µM) or higher (100 µM) concentration of NOC-18. Figure 5 panels A and B show the buildup of product Kyn in the cultures *versus* time under the three conditions for cells expressing IDO1 or TDO. Cells given 5 µM NOC-18 showed greater Kyn buildup than did the control cultures at all time points which resulted in 7 times greater Kyn concentration by 12 h. In contrast, cells given 100 µM NOC-18 generated less Kyn than did the control cells at all time points which resulted in 4 times less Kyn concentration by 12 h. Figure 5 panels C and D report the corresponding levels of ^14^C-heme in IDO1 or TDO after they were immune-precipitated at each time point for the three different culture conditions. In the control cultures the IDO1 and TDO maintained constant levels of ^14^C-heme throughout the time course. Cells given 5 µM NOC-18 stimulated incorporation of ^14^C-heme into IDO1 and TDO that could be detected even after the first hour of exposure, and ^14^C-heme incorporation continued to increase until 6 h of exposure after which it remained at a constant level that was 2 to 3 times above the original ^14^C-heme contents of IDO1 and TDO at time = 0 or that was present in IDO1 or TDO in the control cell cultures. In contrast, in cells given 100 µM NOC-18 there was no gain in ^14^C-heme content in IDO1 or TDO at any time point and instead their ^14^C-heme levels steadily decreased relative to the control cultures, even by the first hour of exposure. Western blot analysis indicated that none of the changes in activity or heme content could be attributed to changes in IDO1 or TDO protein expression in the cells, which remained similar across the time points and different culture conditions (Fig. S9 and S10). Thus, the low NO exposure caused cells to begin steadily incorporating heme into their apo-IDO1 or apo-TDO populations, whereas high NO exposure never stimulated any heme incorporation and instead caused the dioxygenases to begin steadily losing their heme.

**Figure 5.**
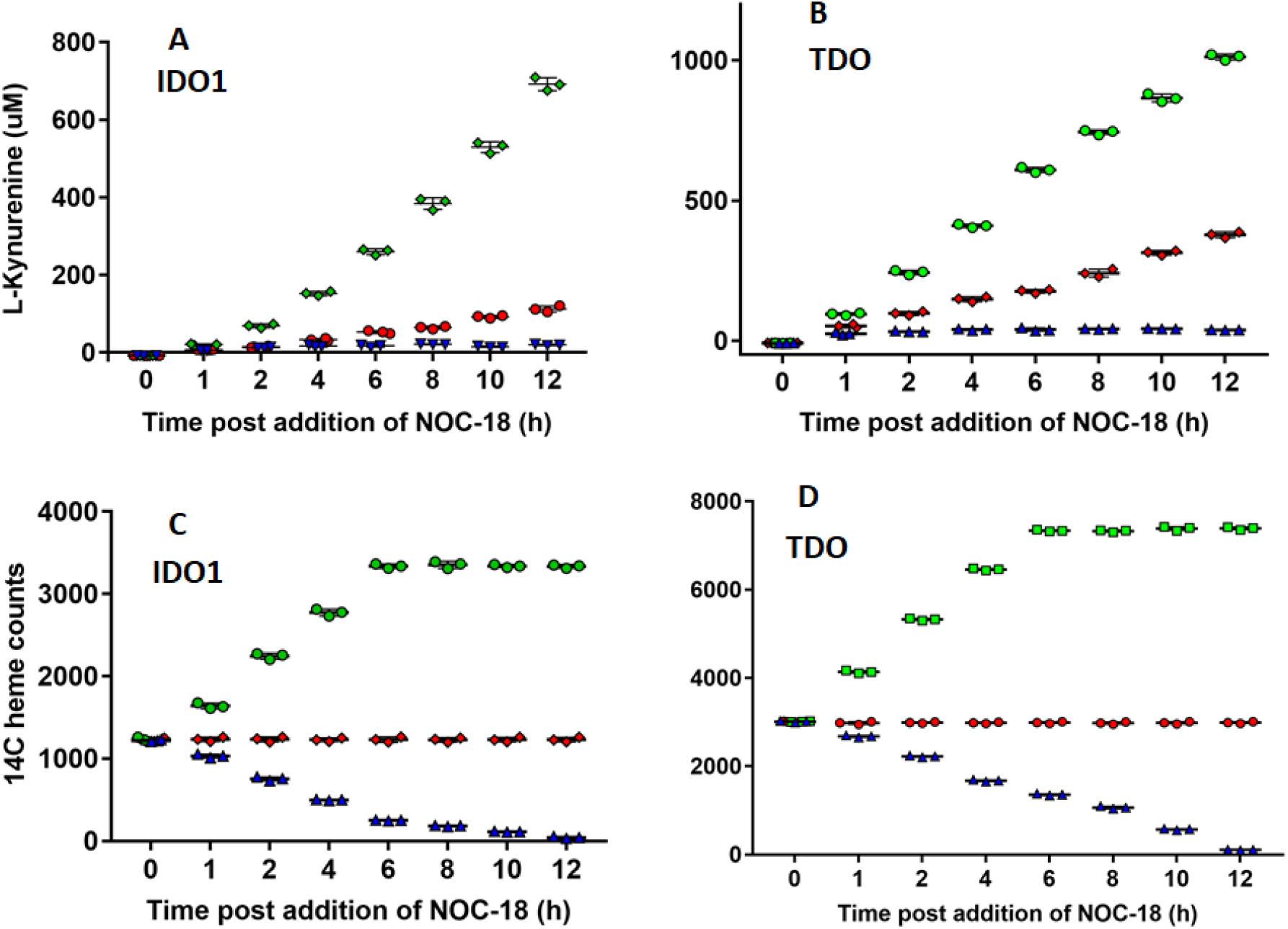
Kinetics of NO-driven changes in IDO1 and TDO activities and heme contents. GlyA-CHO cells were incubated with ^14^C-Gly to produce ^14^C labeled heme and transfected to express IDO1-FLAG or TDO-FLAG. Protein synthesis was halted and NOC-18 at 0 (red), 5 (green), or 100 (blue) µM was added and the cell cultures were analyzed at the indicated times. Panels A and B, Kyn concentrations in the culture media. Panels C and D, ^14^C-heme counts in IDO1-FLAG and TDO-FLAG pull downs. Data are the mean ± s.d.; n=3 experiments. ***p<0.001, ns=not significant, one-way ANOVA.

### NO-stimulates cell heme delivery to IDO1 and TDO through a GAPDH-dependent mechanism

During their normal maturation process, cell heme delivery to apo-IDO1 and apo-TDO requires the formation and participation of a GAPDH-heme complex (29). We therefore investigated if the NO-driven heme acquisitions also involved GAPDH by employing our established siRNA knockdown and rescue strategy (29). GlyA-CHO cells first underwent targeted siRNA knockdown of GAPDH expression or treatment with scrambled siRNA, were given ^14^C-Gly and were then transiently transfected to express IDO1 or TDO alone or in combination with siRNA-resistant forms of wild-type HA-GAPDH or the heme-binding defective HA-GAPDH-H53A variant. Cells were then given 0, 5, or 100 µM NOC-18 and the activities and heme contents of their IDO1 and TDO were determined after 12 h.

The targeted knockdown of GAPDH for 48 h lowered its cell expression by approximately 80% relative to controls, consistent with our previous results (29), the expression of either HA-GAPDH protein in the knockdown cells restored their total GAPDH expression level to a normal value, and the cell IDO1 and TDO expression levels were unaffected by the different treatments (Figs. S11 and S12). Figure 6 panels A and B show that GAPDH knockdown severely diminished Kyn accumulation in the cell cultures expressing IDO1 or TDO, as we reported previously (29), and it also severely diminished the gain in Kyn production that otherwise occurred in response to 5 µM NOC-18. Under both circumstances (without or with 5 µM NOC-18) the expression of the wild-type HA-tagged GAPDH in the knockdown cells restored TDO and IDO1 activities to normal as judged from the levels of Kyn buildup, whereas expression of the GAPDH heme binding mutant (HA-tagged H53A GAPDH) in the knockdown cells did not. In cells undergoing higher NO exposure from 100 µM NOC-18, there was no increase in IDO1 or TDO activities under any circumstance and instead their activities were inhibited relative to the control cells. Figure 6 panels C and D reveal that the GAPDH knockdown and rescue procedures impacted the heme contents of IDO1 and TDO in a manner that matched the impact on their catalytic activities, in both the control cells and in the cells that were treated with 5 µM NOC-18. We conclude that the cell heme allocation into apo-IDO1 and apo-TDO that was driven by 5 µM NOC-18 and their consequent gain in activity is dependent on the cell GAPDH expression level and on the ability of GAPDH to bind intracellular heme.

**Figure 6.**
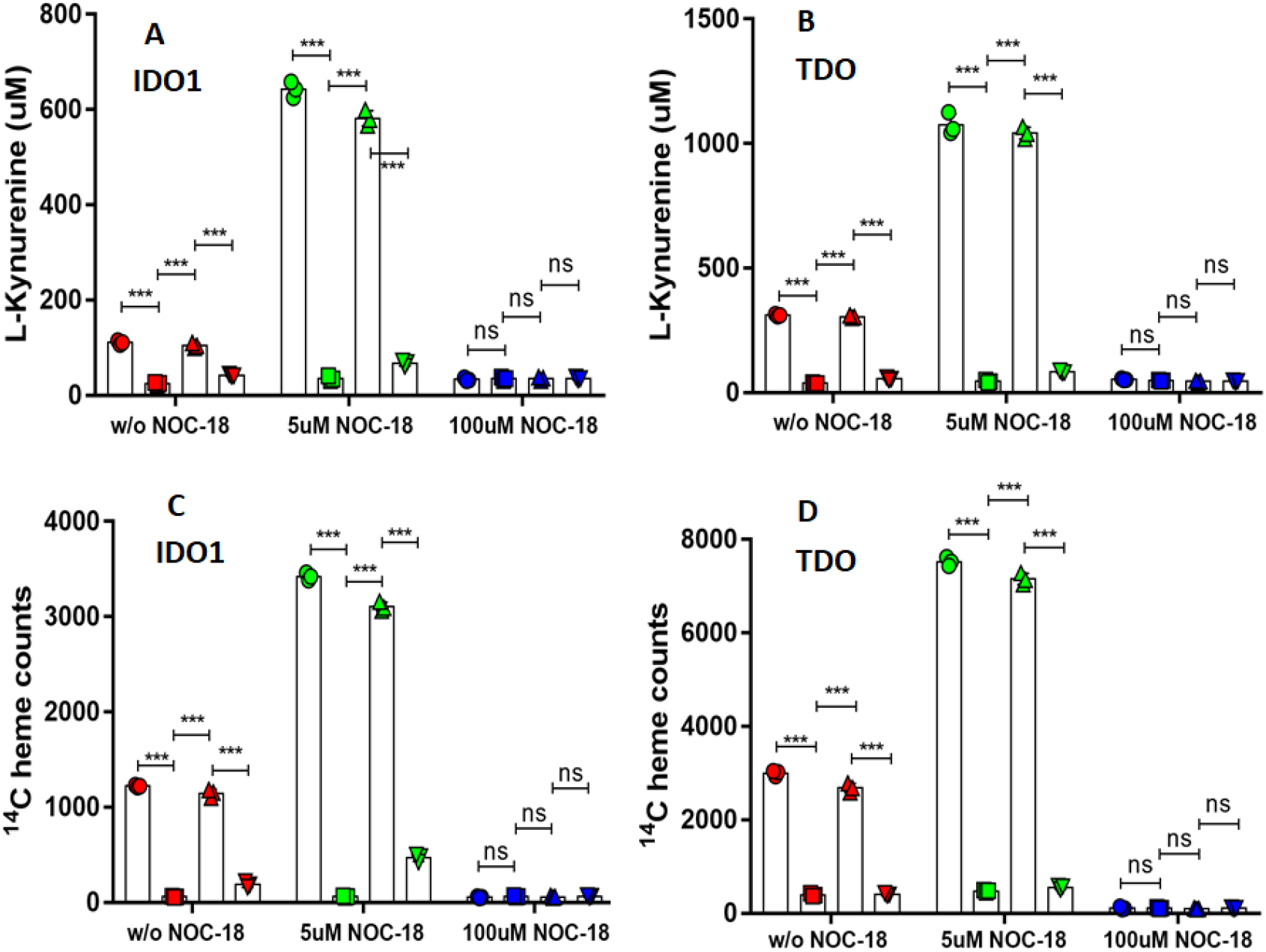
NO-driven cell heme allocation to apo-TDO and apo-IDO1 is GAPDH-dependent. GlyA-CHO underwent siRNA knockdown of GAPDH expression or treatment with scrambled siRNA, were given ^14^C-Gly and then transiently transfected to express IDO1 or TDO alone or in combination with siRNA-resistant forms of wild-type HA-GAPDH or the heme-binding defective HA-GAPDH-H53A variant. Cells were then given 0, 5, or 100 µM NOC-18 and the Kyn production (Panels A and B) and ^14^C-heme contents of IDO1 and TDO (Panels C and D) were determined after 12 h. Circle, control siRNA; square, GAPDH siRNA; triangle, GAPDH siRNA plus HA-GAPDH; inverted triangle, GAPDH siRNA plus H53A HA-GAPDH. Data are the mean ± s.d.; n=3 experiments. ***p<0.001, ns=not significant, one-way ANOVA.

### NO-stimulated heme insertion into IDO1 requires Hsp90

Because Hsp90 is known to drive heme insertion into apo-IDO1 but not into apo-TDO during their normal maturation process (29), we investigated if Hsp90 was also needed for NO-driven heme allocation. We used the specific inhibitor Radicicol to test how blocking the Hsp90 ATPase function in GlyA-CHO cells would impact the NO-driven change in IDO1 and TDO activities and heme contents. Figure 7 panels A and B show that a 6 h pre-incubation with Radicicol completely prevented 5µM NOC-18 from stimulating cells to increase the activity and heme content of IDO1, while in contrast it had no ability to block 5 µM NOC-18 from increasing cell TDO activity and heme content. Western blot analyses showed that Radicicol treatment did not change the expression levels of IDO1 or TDO under any circumstance, which remained constant throughout the experiment (Figs. S13 and S14). These results indicate that functional Hsp90 is needed for the NO-stimulated heme allocation into apo-IDO1 but not into apo-TDO.

**Figure 7.**
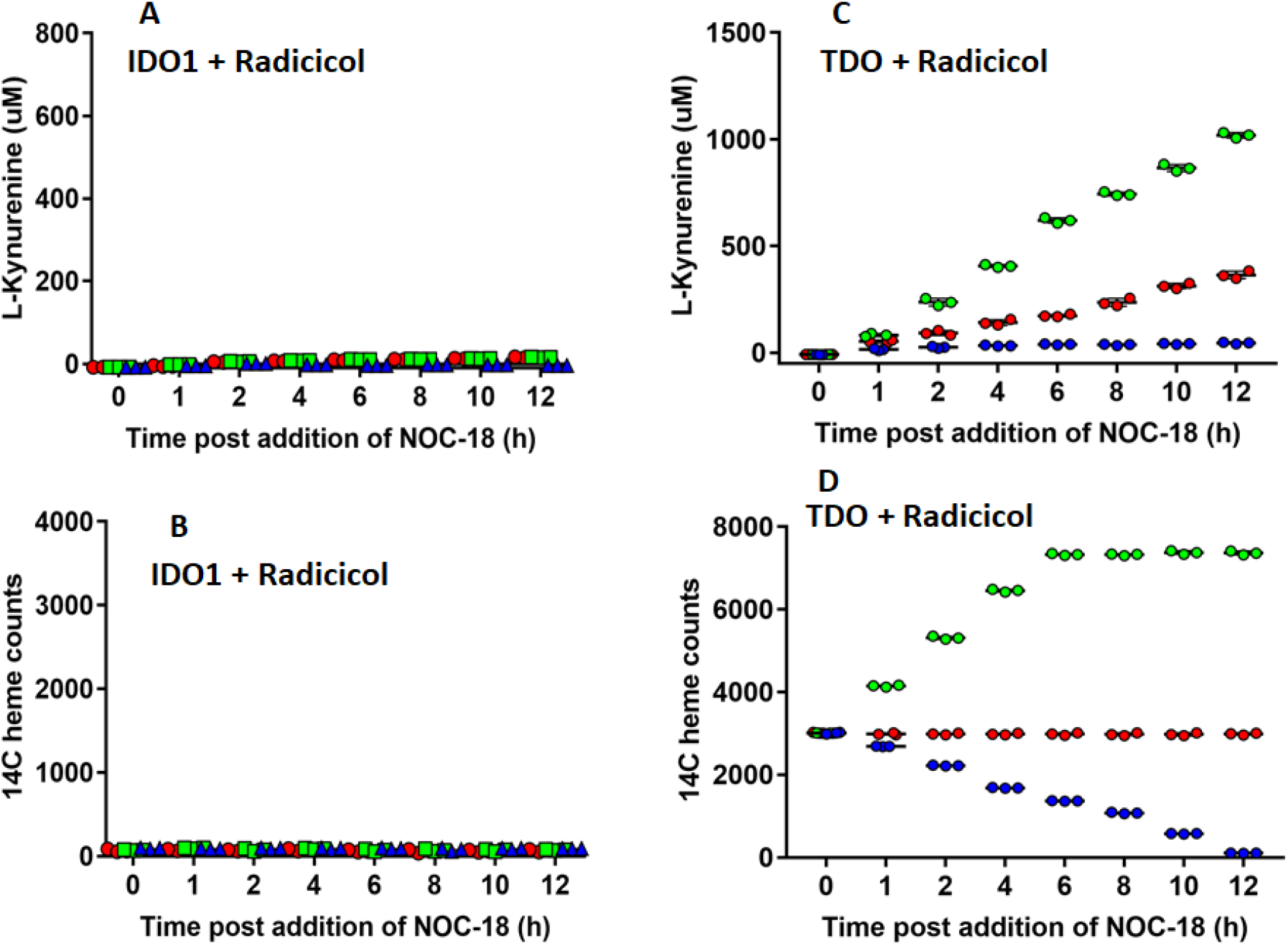
Hsp90 is needed for NO-driven heme allocation to apo-IDO1 but not to apo-TDO. GlyA-CHO cells were given ^14^C-Gly and transfected to express IDO1 or TDO, then given 10 uM Radicicol for a further 6 hrs. NOC-18 at 0, 5 or 100 uM was added to the cells in the continued presence of Radicicol (indicated in red, green, and blue respectively, time = 0) and cells were harvested at the indicated times. Panels A and B, IDO1 Kyn production and ^14^C-heme contents. Panels C and D, TDO Kyn production and ^14^C-heme contents. Data are the mean ± s.d.; n=3 experiments. ***p<0.001, ns=not significant, one-way ANOVA.

## Discussion

We found that NO can positively or negatively regulate IDO1 and TDO activities in cells through its influence on cell heme allocation. Cells exposed to a low-level of NO generation were stimulated to increase the heme contents and catalytic activities of their IDO1 and TDO. Beyond this range, NO steadily lost its positive effect and ultimately caused a loss of the existing heme that was bound within either dioxygenase enzyme. This bimodal effect was observed when NO was released by a chemical NO donor or was released naturally by immune-activated macrophage cells. It was also independent of what cell type expressed IDO1 or TDO and whether the cells naturally expressed the dioxygenases or did so by transfection. Thus, we conclude that NO has a broad ability to both positively and negatively impact cell IDO1 and TDO activities through it regulating cell heme allocation to and from the dioxygenases.

Our experiments utilized cells that were cultured under standard conditions and thus contained their normal or resting levels of heme. In this circumstance both IDO1 and TDO are known to only be partially heme-saturated (29), which allowed us to find that low NO exposures promote heme allocation into the apo-IDO1 and apo-TDO subpopulations that are naturally present in the cells. We saw that NO-driven heme allocation into apo-IDO1 or apo-TDO began within the first hour of the low NO exposure and continued in a steady manner for 6 h until the TDO and IDO1 heme levels had increased by about 3- to 4-fold. The overall effect on their activities was significant and led to a 7-fold increase in the Kyn produced in the cell cultures.

It is remarkable how low of an NO exposure triggered the cells to allocate heme into apo-TDO and apo-IDO1. NOC-18 has the slowest NO release rate of any NO donor that is commercially-available. Despite this, the NO generated by the lowest NOC-18 concentration that we utilized (0.1 µM) still caused a measurable gain in the IDO1 and TDO heme contents over a 12 h period, and exposing the cells to only 5 µM NOC-18 promoted an optimal heme allocation, with an increase in the IDO1 and TDO heme contents being observable even within the first hour exposure. The NO release from NOC-18 at these concentrations is extremely low, 0.13 and 6.6 nM/min respectively for 0.1 µM and 5 µM NOC-18 based on measures we made under our cell culture conditions. In general, NOC-18 has been reported to create solution NO concentrations that are about 1000 times less than its initial concentration (45). This suggests that NO concentrations of only 100 pM to 5 nM were being reached in our cell cultures yet still could stimulate cells to allocate heme into apo-IDO1 and apo-TDO. Such concentrations of NO can enable metal-nitrosyl formation, for example NO binding to the heme in sGC to activate its cGMP synthesis, and also can drive cells to allocate heme into apo-sGCβ (44,46), but are somewhat lower than the NO concentrations required to stimulate cells to phosphorylate ERK or Akt, and are 20 to 80 times lower than the NO concentrations required to promote HIF-1α stabilization or P53 phosphorylation in cells (47). Thus, stimulating cell heme allocation may be one of the more sensitive NO effects in biology. It is equally remarkable how narrow was the window of NOC-18 concentration that could promote cell heme allocation to apo-IDO1 and apo-TDO. Its effectiveness peaked at only 5 µM NOC-18 and then decreased fairly sharply in all four cell types that we examined, such that the positive effect was completely lost by 50 µM NOC-18, which is a concentration that still generates a relatively low NO exposure. Indeed, because NOC-18 has almost never been used below 50 µM in the literature and instead is typically used at concentrations of 100 to 2000 µM, we suspect that the impact of very low NO on other hemeproteins and in biological systems in general has been understudied and to a large part is still overlooked. NO exposure is known to increase the level of exchangeable heme inside cells as detected by an intracellular heme sensor (48). This effect occurred over tens of minutes following NO exposure, which is consistent with the kinetics for the NO-driven heme allocations to apo-TDO and apo-IDO1 that we observed here. NO is also known to speed heme intake and utilization within mammalian cells (49) and NO can trigger heme transfer between purified proteins (50). This suggests that NO may improve heme availability and its mobility inside the cell and deserves further study. We also found that the NO-driven heme allocations to apo-IDO1 and apo-TDO relied on formation of a GAPDH-heme complex in the cells, and for IDO1 also relied on Hsp90 function, which means that the NO acts through a mechanism that involves the same machinery that cells normally use to deliver heme to these dioxygenases (29) and to several other hemeproteins during their maturation (51-54). Within this context NO could act in several ways. The initial heme provision to apo-IDO1 and apo-TDO that occurs within the first hour of low NO exposure likely results from a relatively direct redistribution of the existing cellular heme, for example by NO possibly promoting heme transfer from pre-existing GAPDH-heme complexes in the cell or by it increasing heme loading onto GAPDH. Beyond this time point the possible mechanisms of action can expand and conceivably include NO-induced changes in cell heme biosynthesis, protein expression, or NO-based post-translational protein modifications. This warrants further study. We also note that the heme allocations to apo-IDO1 and apo-TDO induced by low NO exposure did not reverse after reaching a maximal level after 6 h exposure. This suggests there is a range of low NO exposure where negative impacts of NO on heme allocation never manifest or do so in ways that can be compensated for by protective processes that may be operating constitutively within the cells.

Regarding how higher NO exposure negatively impacted TDO and IDO1, we observed it inhibited their activities, consistent with previous reports showing NO inhibits IDO1 activity at higher concentrations (39). Curiously, we found that the higher NO exposures did not trigger any initial heme allocation into apo-IDO1 and apo-TDO and instead only caused the heme-containing IDO1 and TDO populations that were present in the cells to steadily lose their heme. This is reminiscent of reports showing that exposure to higher NO levels prevented cells from allocating heme into globins, NOS enzymes, cytochrome P450s, and catalase (43), and could even result in heme loss from cytochrome P450’s (55). However, it contrasts with recent results we obtained for the hemeprotein sGCβ, where we found that a low or high level of NO exposure both caused cells to allocate heme into apo-sGCβ (44). This apparent discrepancy implies that higher NO levels may impact cell heme allocations differently depending on the identity of the recipient hemeprotein, and merits further investigation.

How NO blocks cell heme allocations or causes heme loss from proteins is still mostly unclear. We know that it blocks cell heme allocation to NOS by causing buildup of a particular post-translational modification in GAPDH, namely Cys-nitrosated GAPDH (SNO-GAPDH) (56). Whether this is a common mechanism whereby NO blocks cell heme allocations to other hemeproteins whose heme deliveries are also GAPDH-dependent (29,53,54,56) is an intriguing possibility. Because cells also express enzymes that denitrosate SNO-GAPDH and recover its function (57), their presence in cells could conceivably diminish or delay the buildup of SNO-GAPDH and thus allow for a window of NO levels to have a “beneficial” effect on GAPDH-dependent heme allocations. In any case, it seems likely that the mechanisms by which low NO promotes cell heme allocations differ from the mechanisms by which high NO blocks heme allocations or even causes heme loss in hemeproteins. Together, these mechanisms likely combine to allow NO to up- and down-regulate IDO1 and TDO activities in cells and can now be further investigated.

NO-driven cell heme allocation into TDO and IDO1 may provide the missing information that can finally explain the classical observation that immune activation in rats caused a temporary increase in their liver TDO activity and heme content, as reported decades ago (11,12,30,36), and may also explain how immune stimulation or NO exposure was found to change IDO1 activity in cells and animals (39,58). Indeed, it is remarkable how the time course of NO-driven ^14^C-heme incorporation into apo-TDO or apo-IDO1 as we showed here (Fig. 5) correlates with the reported gain in liver TDO activity and heme content in rats following an LPS injection (13), which in turn correlates with the reported increase in the blood levels of nitrate (an NO breakdown product) in mice over time following an LPS injection (41). Our finding that NO generated by lower numbers of LPS-stimulated macrophages could increase the activities and heme contents of TDO and IDO1 in neighboring cells recapitulates these early animal studies and overall suggests a mechanism whereby the LPS injections, by inducing NOS expression and an increased NO production in the animals, initially stimulated heme allocation into their apo-TDO (or apo-IDO1) populations, thus boosting their overall activity for Trp metabolism. In short, our current findings potentially explain how biological NO generation can up- or down-regulate Trp metabolism in mammals via an ability to up or down-regulate the heme contents of IDO1 and TDO.

Our findings also have biomedical implications. For example, they may help explain why low-level of NO production has often been found to be pro-cancerous (59,60). Given that IDO1 and TDO are often expressed in malignant tumors and cause unwanted suppression of the immune system via their production of Kyn (27,61), we speculate that a low level of NO might actually be boosting the IDO1 and TDO heme contents, thereby increasing Kyn production and the resultant immune suppression that helps tumor cells escape from immune surveillance. Likewise, a high level of NO via high iNOS expression in tumors is associated with limiting or reversing cancer growth (62). Because IDO1 and TDO both lost their heme in cells given a higher level NO exposure, it is possible that inactivating IDO1 and TDO in this way could occur and thus diminish Kyn production and limit immune suppression, thereby enabling the T-cell differentiation that is needed to achieve tumor surveillance. In auto-immune diseases like severe asthma, high levels of NO are often produced in the airway due to an increased iNOS expression (63). Although IDO1 protein expression is typically up-regulated in asthma, the higher NO levels could cause a heme deficit, thus limiting its Kyn production, diminishing immune suppression, and enabling the lung inflammation that is a characteristic of severe asthma (64). Indeed, IDO1 expressed in airways is reported to have lower than normal activity in both adult and pediatric severe asthma patients (65) and in a mouse asthma model active IDO1 protected the animals from developing severe airway inflammation (58). Taken together, this suggests that the bimodal impact of NO on TDO and IDO1 heme contents and activities as described here is likely to be medically important, and it may present new opportunities to devise strategies to preserve, boost, or limit Kyn production in health and disease.

### Summary

There is a dynamic and bimodal regulation of heme levels in IDO1 and TDO in response to different NO levels by the cell. NO impacts on cell heme allocation could be correlated with prior observations where inflammation and resultant NO generation altered the activities of IDO1 or TDO. Because several other hemeproteins naturally exist in a partially heme-saturated state, and NO also promotes cell heme allocation to sGCβ (44), we speculate that NO may broadly impact the biological functions of hemeproteins in this same bimodal manner.

## Experimental procedures

### 1) Materials

[^14^C]-glycine (0.5 mCi) was purchased from ICN Biomedicals (Cleveland, OH). All other reagents used were purchased from Sigma (St. Louis, MO) unless otherwise mentioned.

### 2) Growth of HepG2 cells

HepG2 cells (ATCC # HB-8065) naturally expressing TDO were cultured in tissue culture treated plates containing EMEM (ATCC # 30-2003) medium containing 10% FBS (Gibco) until 90% confluent after which cycloheximide (Sigma # C7698) was used to treat cells at 5 μg/ml for 12 h to inhibit further protein synthesis. The expression of TDO was checked using anti-TDO antibody (Proteintech # 15880-1-AP). Cells were then utilized for experiments involving treatment with NOC-18 NO donor for indicated doses and time points.

### 3) Plasmids, transfection and expression of human proteins in mammalian cells

Human IDO1-FLAG (Sino Biologicals # HG11650-CF), TDO-FLAG (Sino Biologicals # HG13215-CF), HA-GAPDH-WT and HA-GAPDH-H53A were transfected in HEK293T cells (ATCC # CRL-11268) and GlyA-CHO cells (glycine auxotrophic for growth and a gift from Dr. P. J. Stover, Cornell University) using Lipofectamine 2000 (Invitrogen # 11668019). Briefly, HEK293T cells were grown in DMEM medium containing 10% serum. Post transfection of FLAG-IDO1/TDO plasmids, cells were allowed to express the heme proteins for 48 h after which cycloheximide was used to treat cells at 5 μg/ml for 12 h to inhibit further protein synthesis. Cells were then utilized for experiments involving treatment with NOC-18 for indicated doses and time points. GlyA-CHO cells were grown in DMEM/F-12 medium without Glycine (Caisson Labs # DFP04) with 10% heme depleted serum and 2 μCi/ml of [^14^C] glycine for 72 h post transfection of FLAG-IDO1/TDO plasmids. At this stage the heme proteins are expressed and have a basal level of ^14^C labeled heme incorporated into them. Cycloheximide was used to treat cells at 5 μg/ml for 12 h to inhibit further protein synthesis. Cells were then utilized for experiments involving treatment with NOC-18 for indicated doses and time points.

### 4) Treatment of cells with NOC-18

NOC-18 (Dojindo Molecular Technologies # N379-12) was freshly dissolved in cell culture grade sterile PBS at a stock concentration of 10 mM and added to phenol red free cell culture medium containing 10% serum and L-Trp at 2 mM final concentration. This medium was used to treat the cells for the indicated time points. The pH of the medium was measured to be 7.0; the reported half-life of NOC-18 at pH 7 at 37 °C was 13 h (Dojindo Molecular Technologies Inc. website for NOC-18).

### 5) Treatment of cells with Radicicol

Hsp90 inhibitor Radicicol (Sigma # R2146) was used to pre-treat cells for 6 h before treatment with NOC-18 at a final concentration of 10 μM. After pre-treatment, cells were maintained in presence of 10 μM Radicicol during NOC-18 treatment for the indicated time points and doses.

### 6) ^14^C labeled heme production in cells and measuring radiolabeled heme counts

^14^C glycine uptake by GlyA-CHO cells generated ^14^C labeled heme which incorporated into FLAG-IDO1/TDO. We maintained a negative control where GlyA-CHO cells were treated with heme synthesis inhibitor succinyl acetone (Sigma # D1415) at 400 μM for 72 h post transfection of FLAG-IDO1/TDO plasmids along with 2 μCi/ml of [^14^C] glycine. In this circumstance the heme proteins are expressed with ^14^C glycine incorporated into the poly-peptide but do not have any heme incorporated into them. The ^14^C counts of the FLAG-IDO1/TDO from these negative controls were subtracted from the experimental samples. Cycloheximide was used to treat cells at 5 μg/ml for 12 h to inhibit further protein synthesis. The method for measuring ^14^C heme counts using a scintillation counter is described in (66).

### 7) Transfection of siRNA and gene silencing

Cell GAPDH protein expression was reduced using siRNA against human GAPDH mRNA. Commercially available siRNA against human GAPDH (# D-001830-01-05) and scrambled siRNA (# D-001810-10-05) were purchased from Dharmacon and used at a final concentration of 100 nM in cultures of mammalian cells of low passage number along with Lipofectamine 2000. The siRNA treated cells were cultured for 72 h before they received transfections with protein expression plasmids as described above.

### 8) IDO1 and TDO activity assay

The enzyme activity of IDO1 and TDO was measured using a colorimetric assay. The cells after treatment with cycloheximide were treated with IDO1/TDO substrate L-Trp at 2 mM in phenol red free DMEM/EMEM medium containing appropriate type of serum (normal or heme depleted) for indicated time to allow for substrate utilization and L-Kynurenine product formation. The medium was collected and de-proteinized by adding an equal volume of 3% trichloroacetic acid (TCA) and incubated at 50 °C for 30 min. The tubes were centrifuged at 9,000 g for 10 min at room temperature to precipitate proteins from the medium. Equal volumes of deproteinized sample were mixed with a freshly made 20 mg/ml solution of p-dimethyl-amino-benzaldehyde (Ehrlich’s reagent; Sigma # 109762) in glacial acetic acid at room temperature to allow for the formation of a yellow colored product. The end point absorbance of this product was measured at 492 nm (Molecular Devices) to determine the concentration of L-Kyn in the medium. A standard curve was obtained using commercial L-Kyn (Sigma # K8625) dissolved in 0.5 N hydrochloric acid in various concentrations using the exact same method.

### 9) Immune-precipitation (IP) and western blot

GlyA-CHO cells were lysed using 50 mM Tris-HCl pH 7.4 buffer with 0.1% Triton X-100, 5 mM Na-molybdate and EDTA-free protease inhibitor cocktail (Roche). Protein concentration was measured using the Bradford method (Bio-Rad # 500-0006). IP pull-downs were performed using 1 mg of whole cell extracts with anti-FLAG antibody (Sigma # F1804). Protein G agarose beads (Millipore # 16-201) were used to pull down the antibody-protein complex. The beads were washed well with lysis buffer, the 1.5 ml tubes were inserted into 5 ml scintillation vials and 4 ml of scintillation fluid (Liquiscint, National Diagnostics # LS-121) were added to each vial. The method for measuring ^14^C heme counts using a scintillation counter is described in (66).

For Western blots to check protein expressions across all samples in all experiments, we lysed cells using 50 mM Tris-HCl pH 7.4 buffer with 0.1% Triton X-100, 5 mM Na-molybdate and EDTA-free protease inhibitor cocktail (Roche). Protein concentration was measured using the Bradford method (Bio-Rad # 500-0006). In each sample 30 μg of cell extract proteins were boiled in Laemmli buffer, resolved onto 10% SDS-PAGE and transferred to PVDF membrane (Bio-Rad # 1620177) and probed for proteins of interest. Western blot was performed with anti-FLAG (Sigma # F1804; dilution 1:1000), anti-GAPDH (Santa Cruz Biotechnology # sc-32233; dilution 1:2500), anti-TDO (Proteintech # 15880-1-AP; dilution 1:1000), anti-iNOS (R & D systems # MAB9502; dilution 1:1000) and anti-α-Tubulin (Santa Cruz Biotechnology # sc-5286; dilution 1:2500). The proteins were detected using chemiluminescence using HRP conjugated secondary antibodies of either anti-mouse (Bio-Rad # 170-6516, dilution 1:10,000) or anti-rabbit (GE Healthcare # NA9340, dilution 1:10,000) origin and ECL substrate (Thermo Scientific # 32106). The images were acquired using a chemidoc system from Bio-Rad.

### 10) Trans-well co-culture experiment

We used a 6-well plate format (Corning # CLS3450) for our transwell co-culture experiments. We seeded RAW264.7 (ATCC # TIB-71) cells in the upper chamber. These cells at a final confluency reached 25% and 100% when they were activated with LPS at 1 μg/ml (Sigma # L2630) for 6h. The bottom chamber had GlyA-CHO cells expressing FLAG-IDO1/TDO in ^14^C glycine containing medium. After initial activation of the RAW264.7 with LPS for 6h, the upper chamber baskets were introduced to the GlyA-CHO cells along with fresh medium in the bottom chamber for 6h. The medium was phenol red free and was used to measure the activities of IDO1/TDO as described previously. Also, nitrite accumulation in this medium was measured using the Griess reagent system (Promega # G2930). The lysates of the RAW264.7 cells were used to check iNOS expression by Western blot. The lysates of the GlyA-CHO cells were used in IP to measure ^14^C heme counts and to determine FLAG-IDO1/TDO expressions by Western blot.

### 11) NO release from NOC-18

The NO mediated conversion of oxy-hemoglobin to met-hemoglobin was used to determine the rate of NO release from NOC-18 at 37 °C. Various concentrations of NOC-18 were added to cuvettes that contained phenol red free DMEM, 10% FBS, 2 mM L-Trp and 10 μM oxy-hemoglobin. The absorbance gain at 401 nm was recorded per minute over a 3h period for each concentration of NOC-18. The rate of NO release was calculated using the difference extinction coefficient of 38 mM^-1^cm^-1^ (43).

### 12) Statistical analyses

All experiments were done in three independent trials, with three replicates per trial. The results are presented as the mean of the three trial values ± standard deviation. The statistical test used to measure significance (p-values) was one-way ANOVA in the software Graph Pad Prism (v9).

## Data availability

All data are contained within the manuscript.

## Supporting information

This article contains supporting information.

## Acknowledgements

We thank members of the Stuehr lab for helpful discussion, and Mr. Joseph Palazzo for excellent technical assistance.

## Abbreviations

Hsp90: heat shock protein 90
sGC: soluble guanylate cyclase
HD: heme depleted
NO: nitric oxide
NOC-18: 2,2ʹ-(Hydroxynitrosohydrazino)bis-ethanamine
DMEM: Dulbecco’s Modified Eagle Medium
PBS: Phosphate-Buffered Saline
IDO1: Indoleamine-2, 3-dioxygenase
TDO: Tryptophan-2, 3-dioxygenase
Trp: Tryptophan; IFNγ, interferon-γ
LPS: lipopolysaccharide
TNF: tumor necrosis factor
Kyn: Kynurenine
GAPDH: Glyceraldehyde 3-phosphate dehydrogenase
NOS: nitric oxide synthase
siRNA: Small interfering Ribo Nucleic Acid
HA: hemagglutinin
CHO: Chinese hamster ovary
ATP: Adenosine triphosphate
cGMP: Cyclic guanosine monophosphate
ERK: extracellular signal-regulated kinase
HIF: Hypoxia-Inducible Factor
EMEM: Eagle’s minimum essential medium
TCA: trichloroacetic acid
EDTA: Ethylene diamine tetraacetic acid
IP: Immune-precipitation
SDS-PAGE: Sodium dodecyl-sulfate polyacrylamide gel electrophoresis
PVDF: Polyvinylidene difluoride
HRP: horseradish peroxidase
ECL: Enhanced chemiluminescence
FBS: Fetal bovine serum
ANOVA: Analysis of variance
Ab: antibody.

