## Supporting Information for "Indoleamine Dioxygenase and Tryptophan Dioxygenase Activities are Regulated through Control of Cell Heme Allocation by Nitric Oxide"


| **Figure number** | **Page number** |
|---|---|
| S1 | S-2 |
| S2 | S-3 |
| S3 | S-4 |
| S4 | S-5 |
| S5 | S-6 |
| S6 | S-7 |
| S7 | S-8 |
| S8 | S-9 |
| S9 | S-10 |
| S10 | S-11 |
| S11 | S-12 |
| S12 | S-13 |
| S13 | S-14 |
| S14 | S-15 |

Fig S1: Rate of NO release from NOC-18 under our experimental conditions.

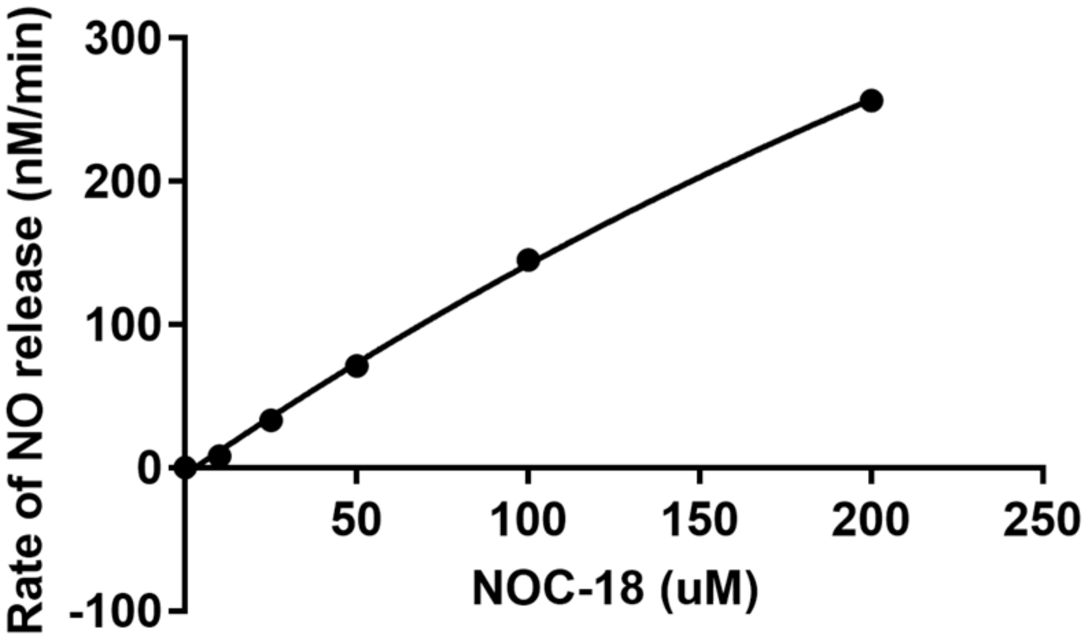

Fig S2: TDO expressions in different NOC-18 treated conditions in HepG2 cells. WB corresponds to Fig. 1.

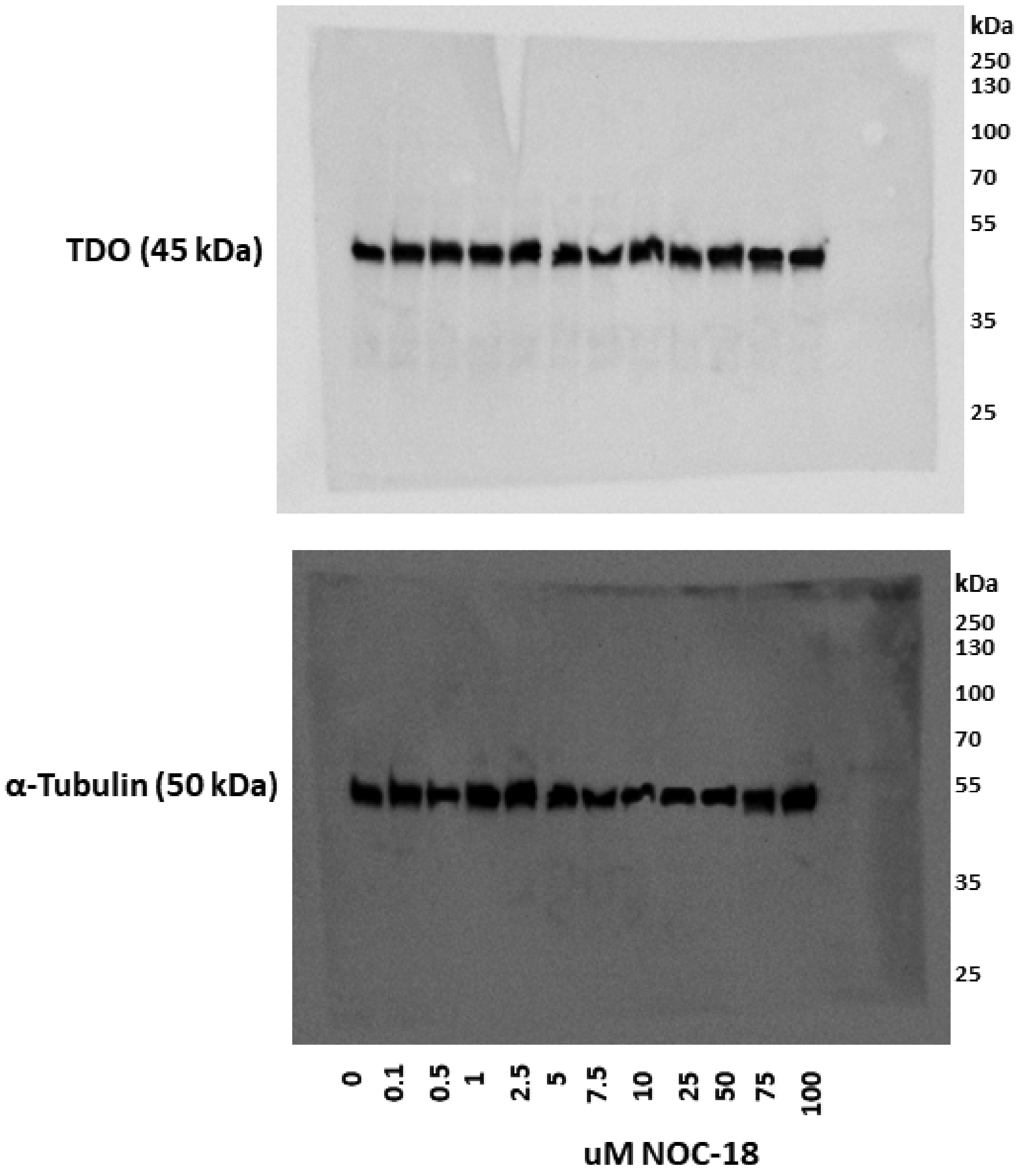

Fig S3: TDO-FLAG expressions in different NO donor treated conditions in HEK293T cells. WB corresponds to Fig. 1.

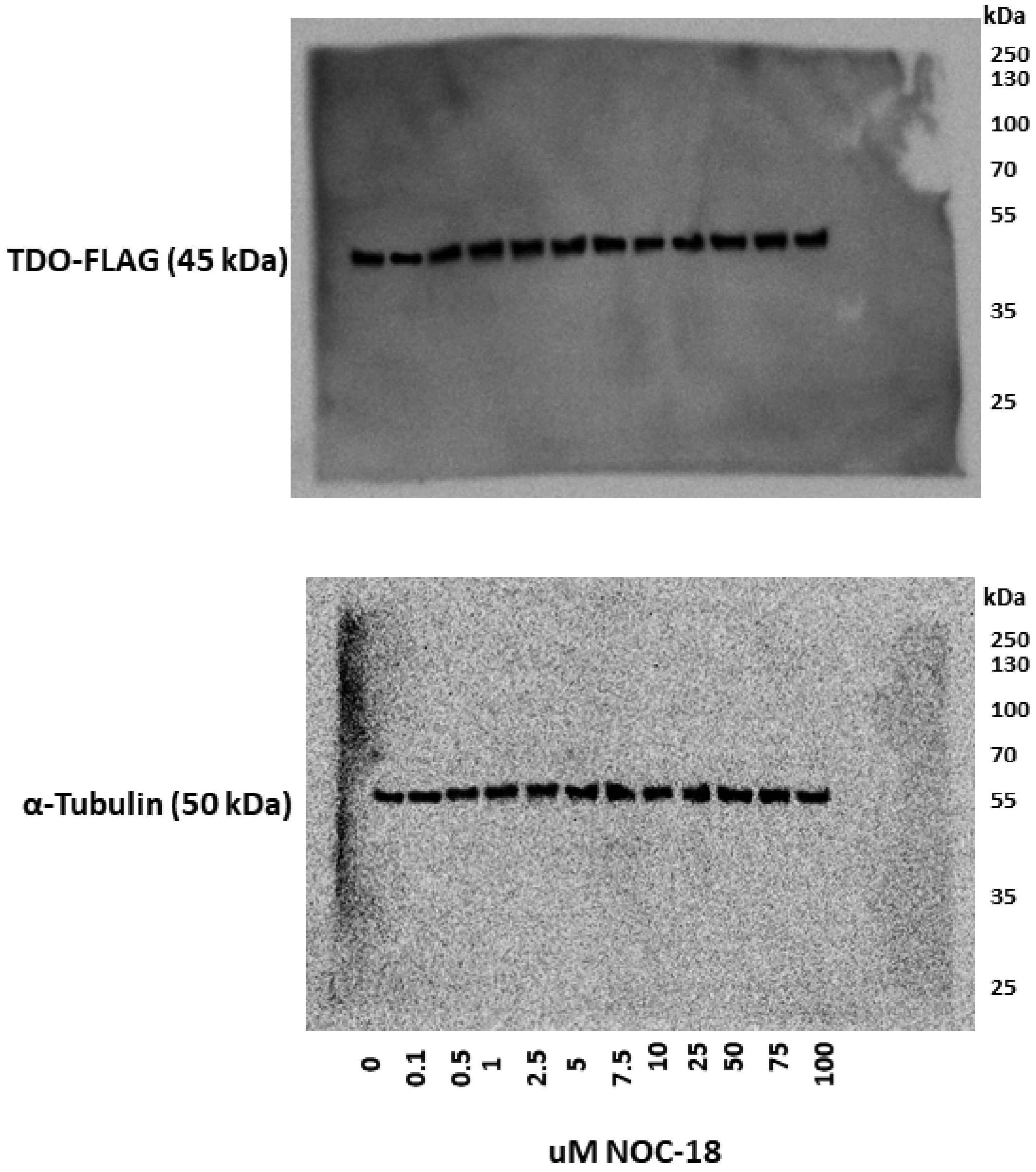

Fig S4: IDO1-FLAG expressions in different NOC-18 treated conditions in HEK293T cells. WB corresponds to Fig. 1.

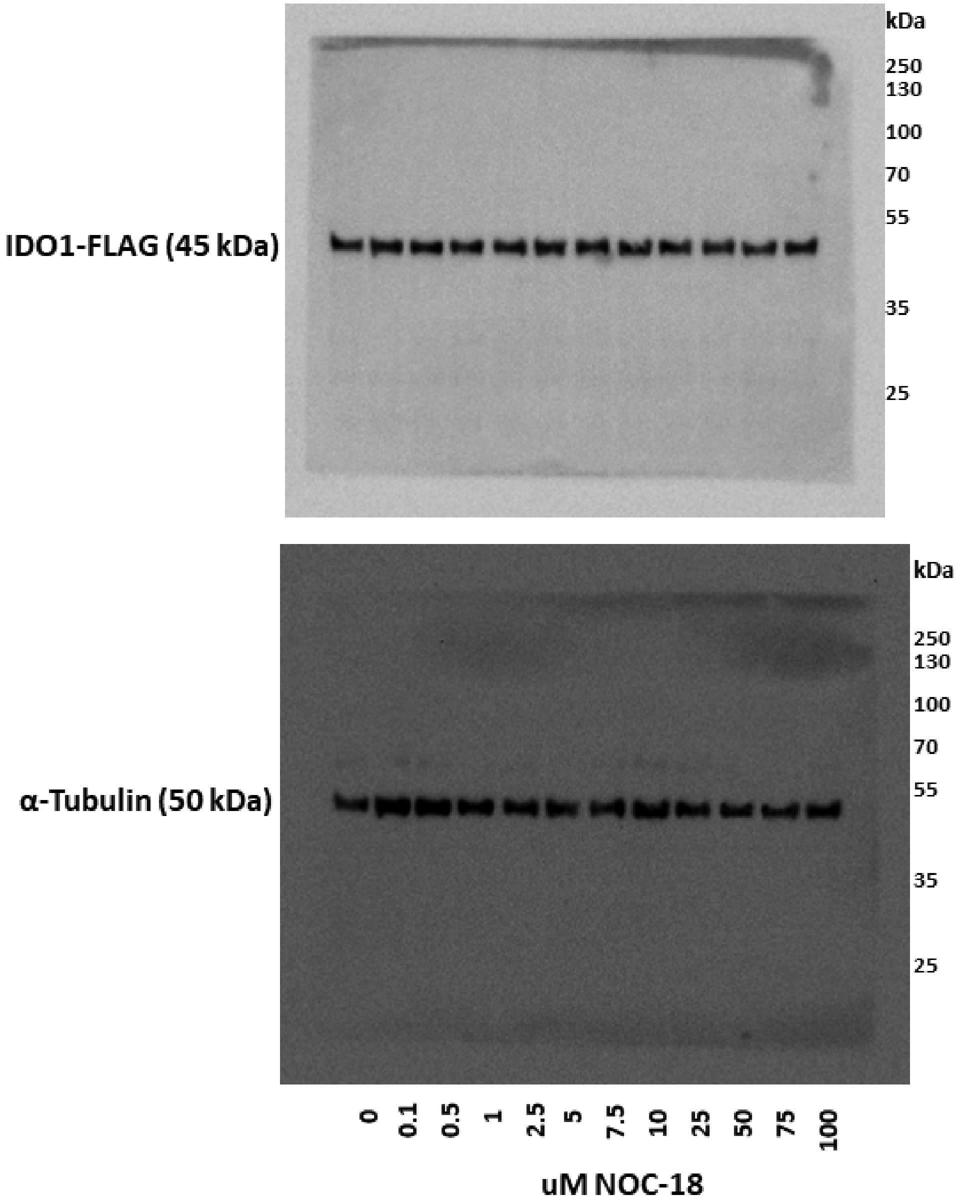

Fig S5: TDO-FLAG expressions in different NOC-18 treated conditions in GlyA-CHO cells. WB corresponds to Fig. 2.

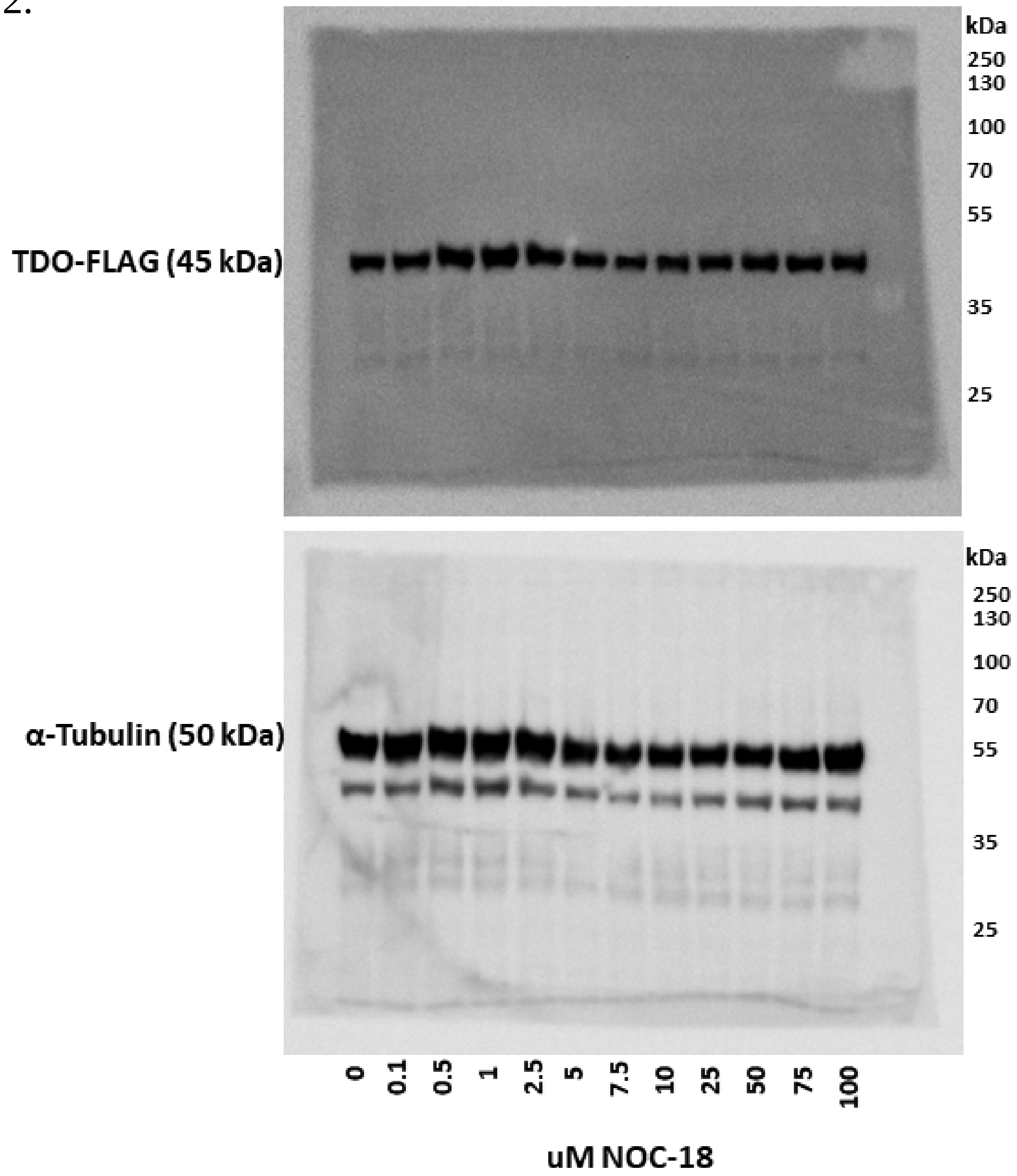

Fig S6: IDO1-FLAG expressions in different NOC-18 treated conditions in GlyA-CHO cells. WB corresponds to Fig. 2.

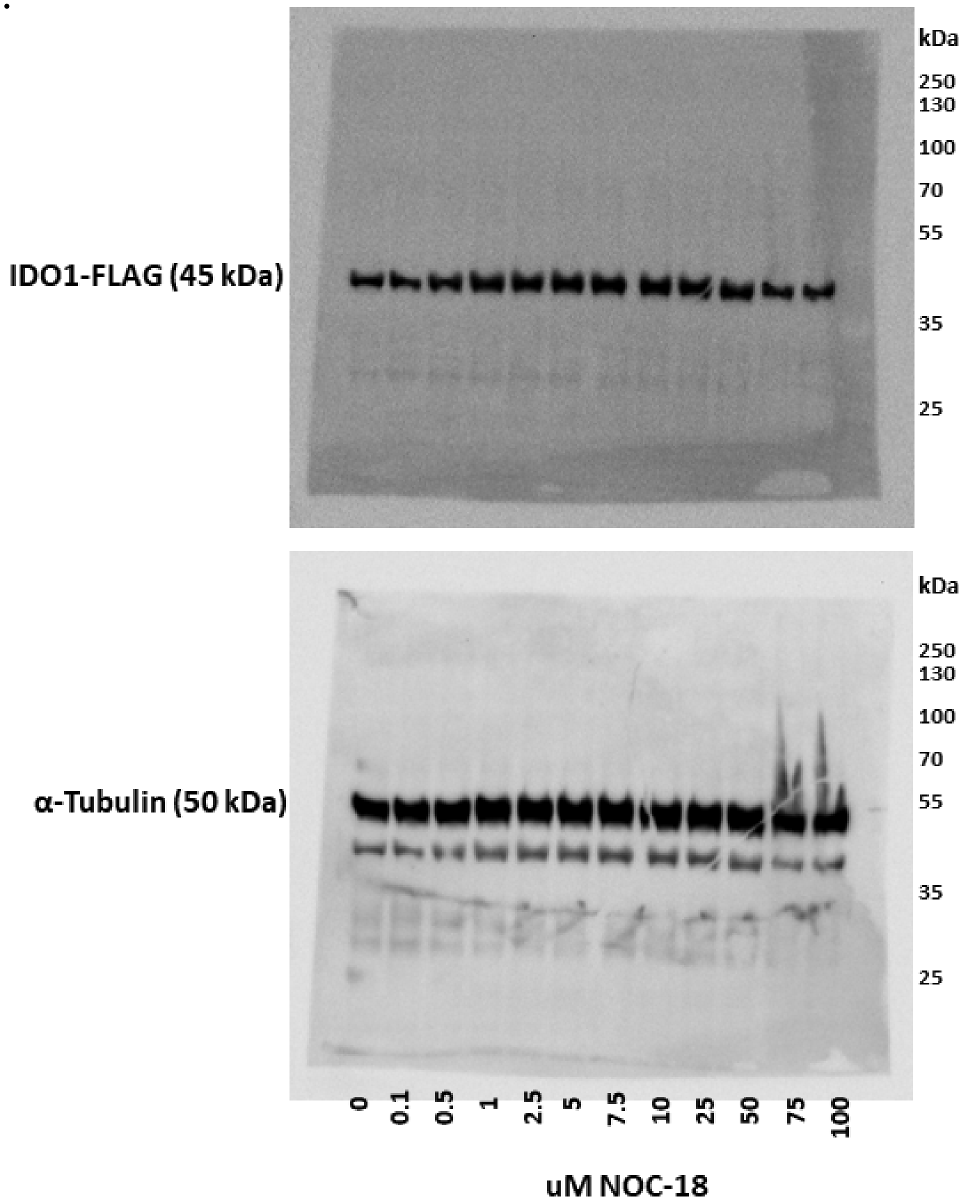

Fig S7. IDO1 and TDO activity and $^{14}C$ heme content in presence of non-activated RAW cells in trans-well co-culture experiment.

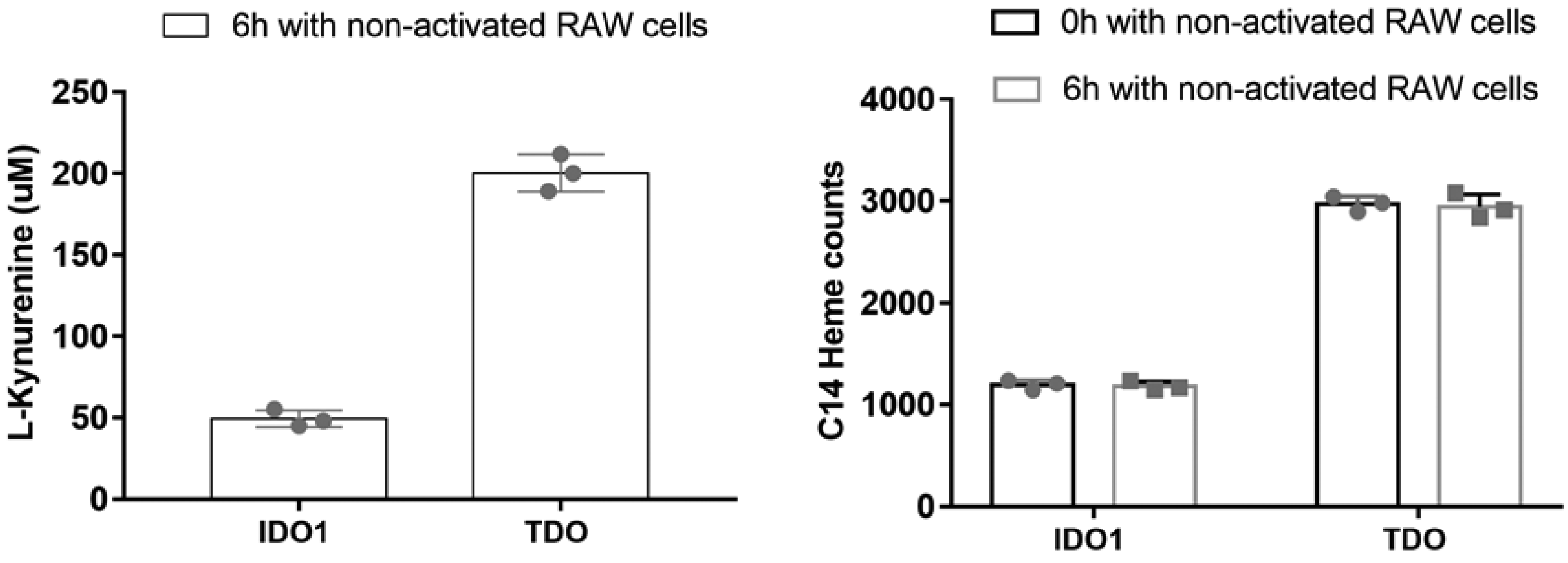

Fig S8: Expressions of IDO1-FLAG and TDO-FLAG in GlyA-CHO cells in the bottom chamber in the trans-well co-culture experiments with RAW264.7 cells in the upper chamber. RAW264.7 cells expression levels of iNOS upon activation with 1 μg/ml LPS for 6h. The WB corresponds to Figs. 3 and 4.

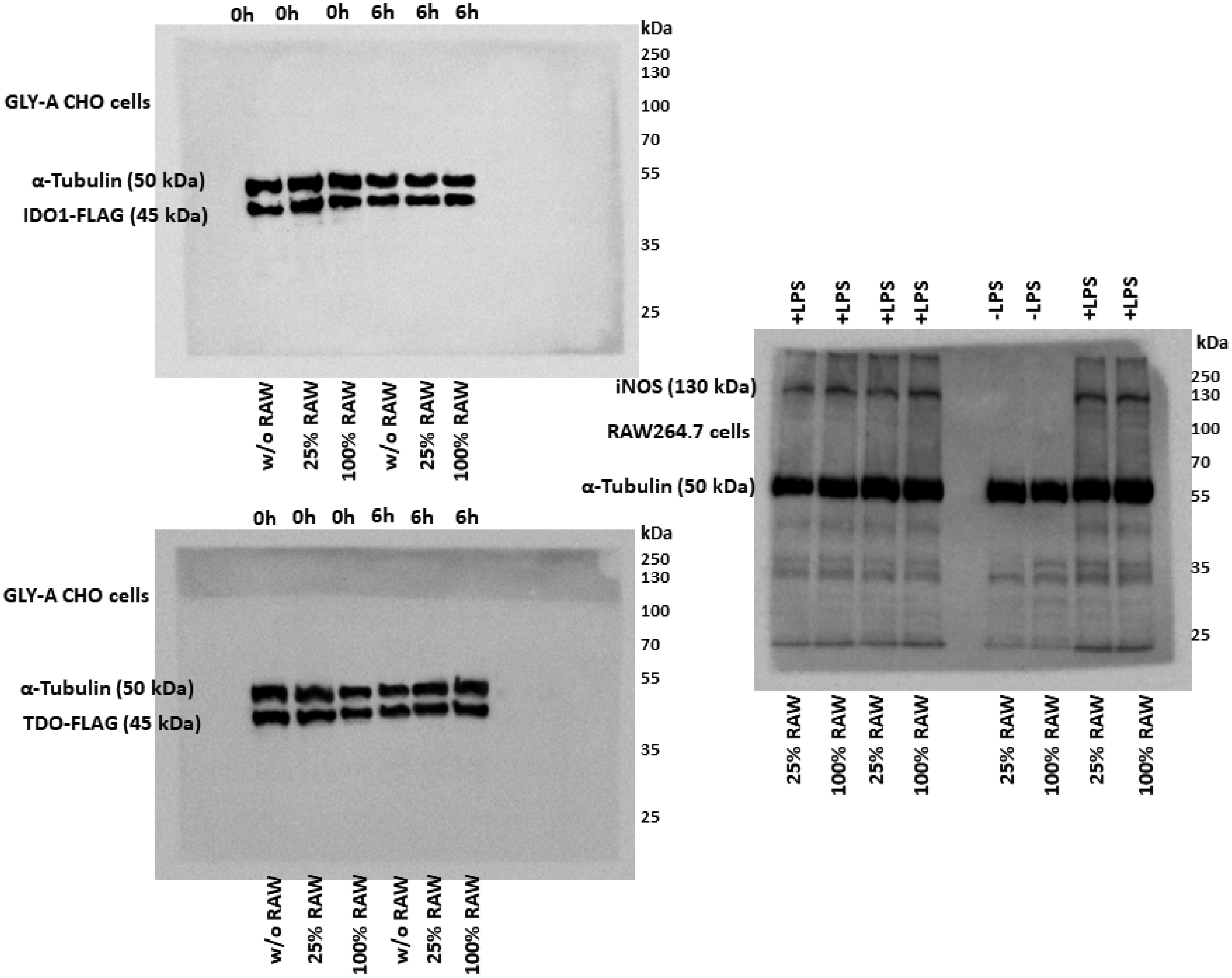

Fig S9: IDO1-FLAG expression levels in GlyA-CHO cells treated without, 5 µM and 100 µM NOC-18 for different time points as mentioned. WB corresponds to Fig. 5.

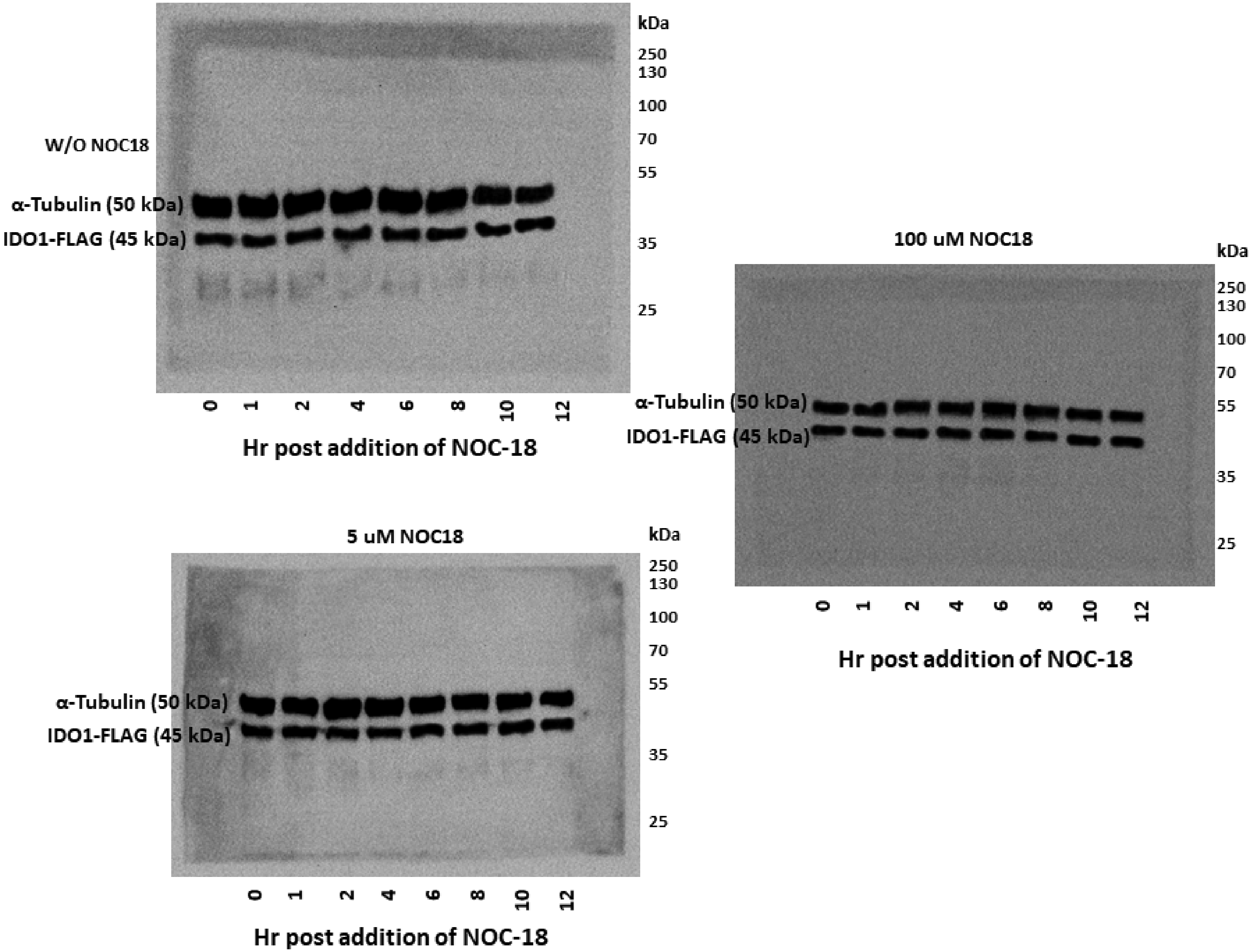

Fig S10: TDO-FLAG expressions in GlyA-CHO cells treated without, 5 μM and 100 μM NOC-18 for different time points as mentioned. WB corresponds to Fig. 5.

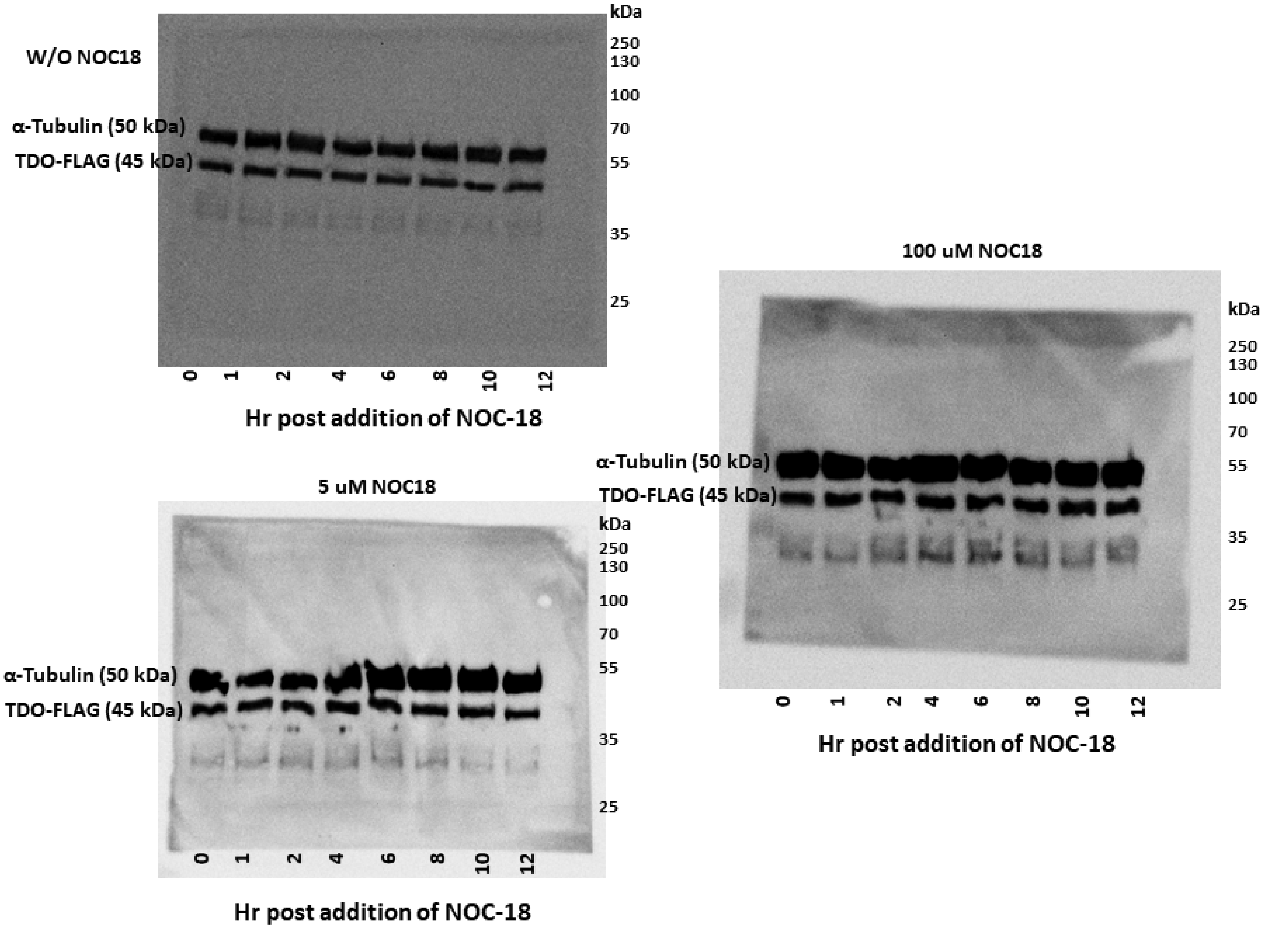

Fig S11: IDO1-FLAG expression in GlyA-CHO cells upon silencing of GAPDH with siRNA and complemented with either WT or H53A HA-GAPDH expression from transfected plasmid. GAPDH was effectively silenced as seen in the blot. WB corresponds to Fig. 6.

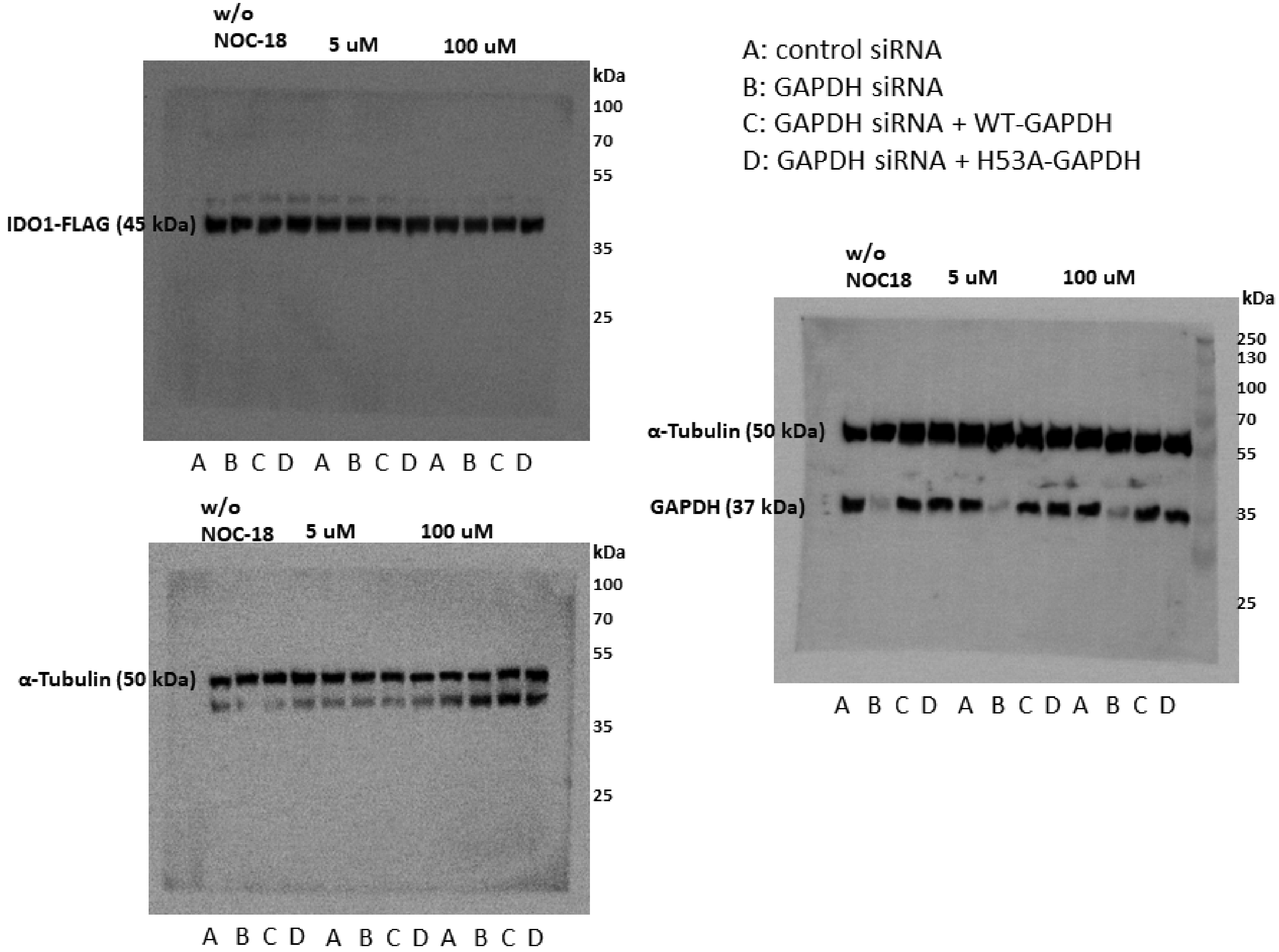

Fig S12: TDO-FLAG expression in GlyA-CHO cells upon silencing of GAPDH with siRNA and complemented with either WT or H53A HA-GAPDH expression from transfected plasmid. GAPDH was effectively silenced as seen in the blot. WB corresponds to Fig. 6.

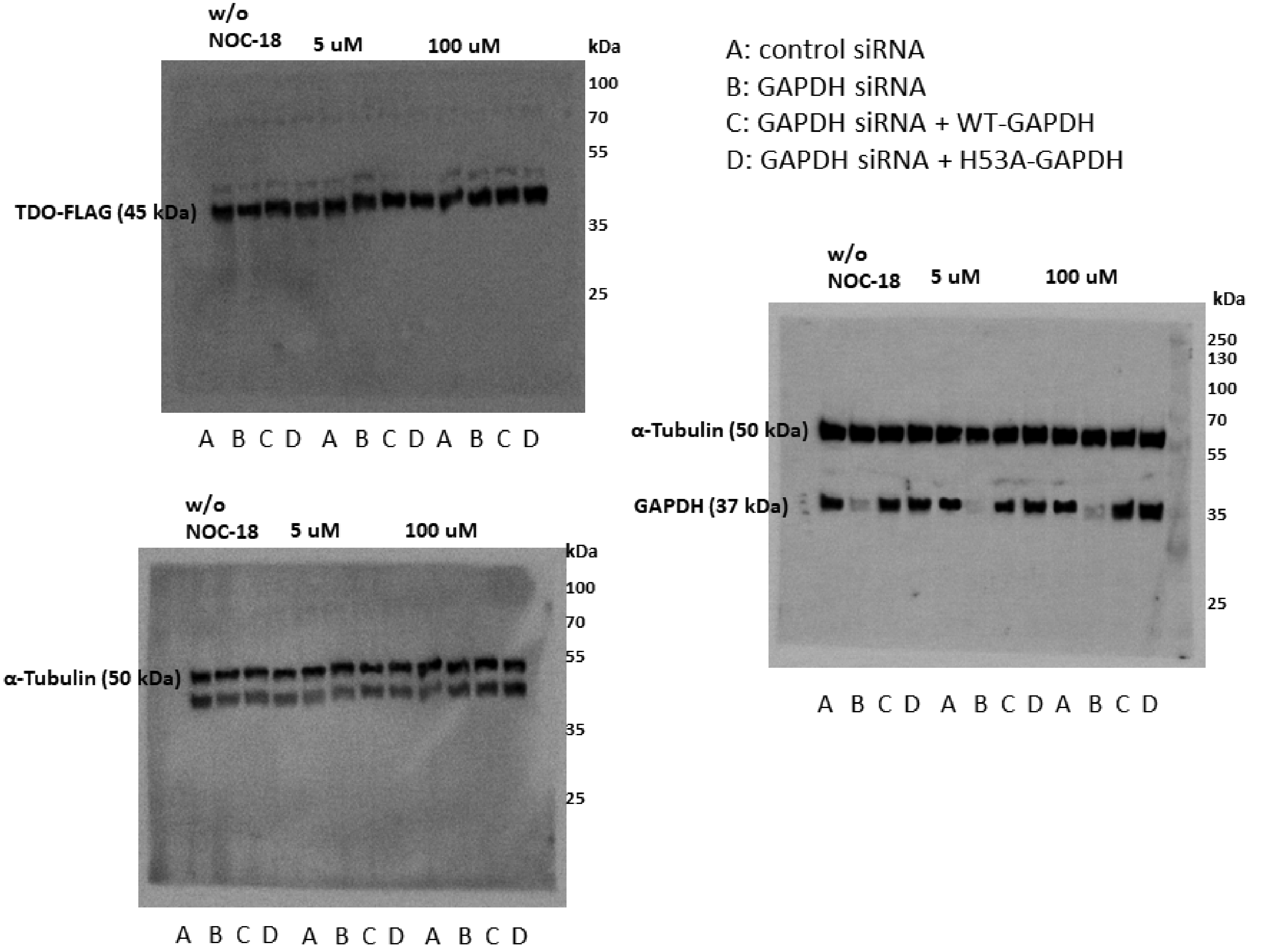

Fig. S13. IDO1-FLAG expression in GlyA-CHO cells upon pre-treatment with 10 μM Radicicol for 6h followed by treatment with different doses of NOC-18 for the indicated time points in presence of Radicicol. WB corresponds to Fig. 7.

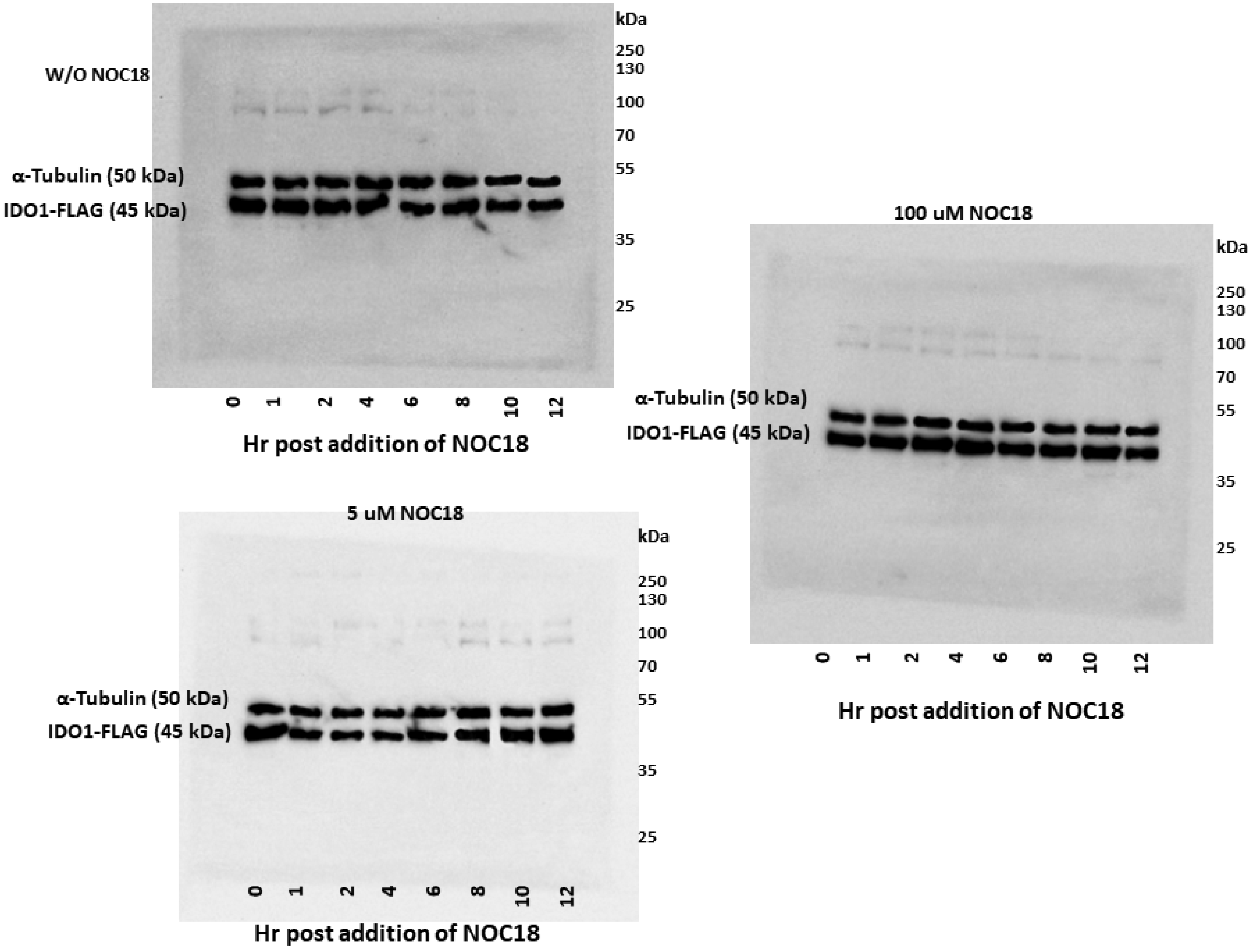

Fig. S14. TDO-FLAG expression in GlyA-CHO cells upon pre-treatment with 10 µM Radicicol for 6h followed by treatment with different doses of NOC-18 for the indicated time points in presence of Radicicol. WB corresponds to Fig. 7.

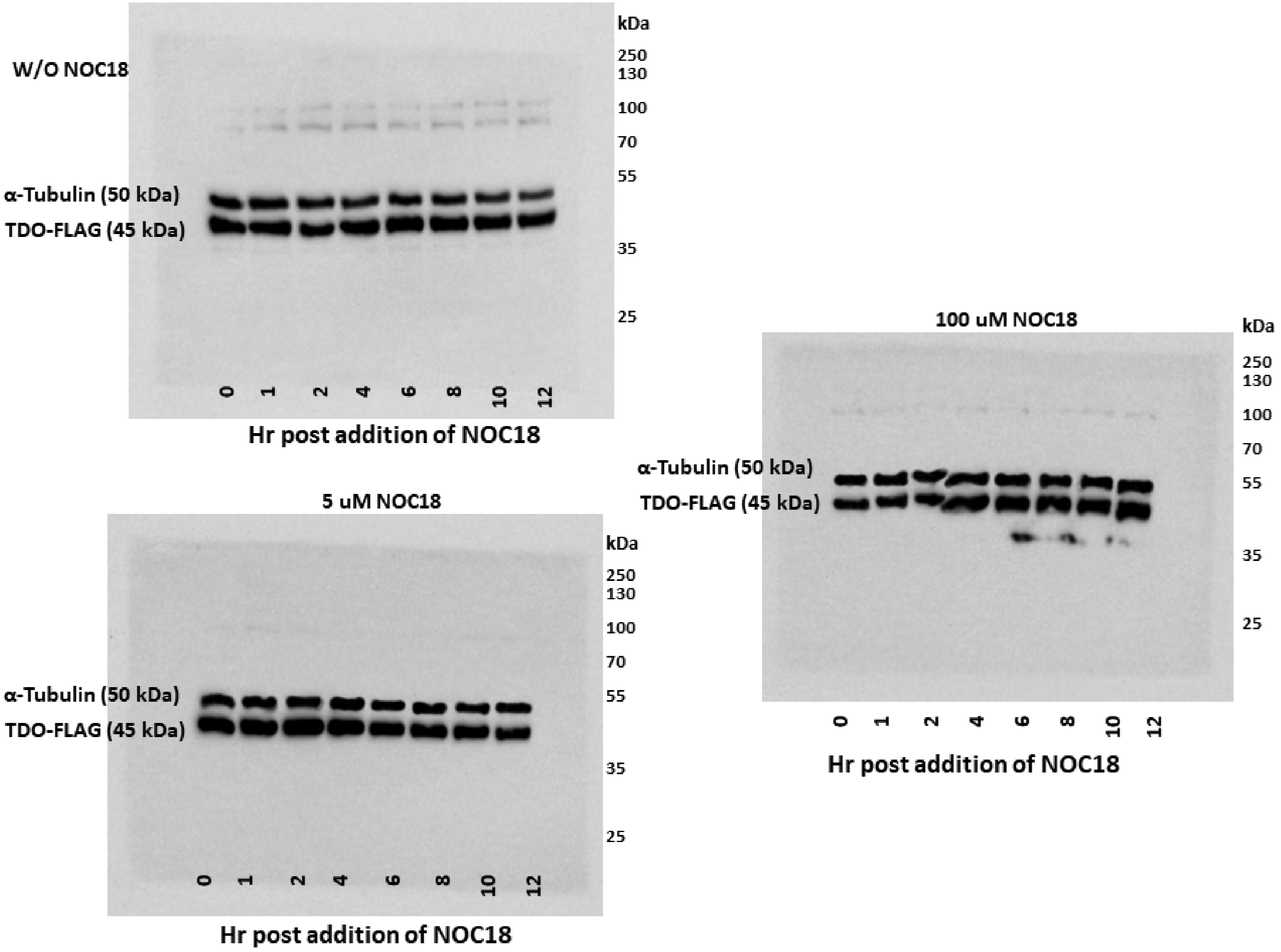
